# Spontaneous activity of striatal projection neurons supports maturation of striatal inputs to substantia nigra dopaminergic neurons

**DOI:** 10.1101/2024.01.05.574371

**Authors:** Bojana Kokinovic, Patricia Seja, Angelica Donati, Maria Ryazantseva, Alban de Kerchove d’Exaerde, Serge N. Schiffmann, Tomi Taira, Svetlana M. Molchanova

**Author notes:** Authors contributed equally to the work (Bojana Kokinovic, Patricia Seja).

## Abstract

Spontaneous activity of neurons during early ontogenesis is instrumental for stabilization and refinement of developing neuronal connections. The role of spontaneous activity in synaptic development has been described in detail for cortical-like structures. Yet, very little is known about activity-dependent development of long-range inhibitory projections, such as projections from striatum. Here, we show that striatal projection neurons (SPNs) in dorsal striatum are spontaneously active in P4-P14 mice. Spontaneous activity was detected in both direct-pathway SPNs (dSPNs) and indirect-pathway SPNs (iSPNs). Most of the spontaneously active cells were in striosomes – a chemical compartment in striatum defined by expression of µ-opioid receptor. Higher excitability of both striosomal dSPNs and iSPNs was related to their intrinsic excitability properties (higher action potential half-width and IV slope). Tonic activation of muscarinic M1 receptor maintains the spontaneous activity of striosomal SPNs, the effect being stronger in iSPNs and weaker in dSPNs. To investigate if the neonatal spontaneous activity is needed for the stabilization of SPN long-range projections, we chemogenetically inhibited striosomal SPNs in neonatal animals and studied the efficiency of striatonigral projections in adult animals. Inhibition of striosomal SPNs by chronic CNO administration to P6-14 pups caused a reduction in the functional GABAergic innervation and in the density of gephyrin puncta in dopaminergic neurons of substantia nigra pars compacta of the adult (P52-79) animals. Chronic administration of CNO later in development (P21-29), on the contrary, resulted in higher mIPSC frequency in dopaminergic cells of the adult animals. Thus, the activity-dependent stabilization of striosomal projections has different developmental phases, and the long-term outcome of perturbations in these processes depends on the developmental period when they occur. Taken together, our results demonstrate that spontaneous activity of SPNs is essential for the maturation and stabilization of striatal efferents.

## Introduction

During early stages of brain development, when sensory inputs are not yet fully functional, maturation of the neuronal networks is guided by the spontaneous activity, generated by the developing neurons. Activity-dependent development is described in detail for the brain areas, where projection neurons are glutamatergic (e.g. cerebral cortex (Molnár et al., 2020), thalamus (Martini et al., 2021) and hippocampus (Cossart & Khazipov, 2022; Huupponen et al., 2013)). However, maturation of the neuronal networks in the subcortical regions interconnected with the inhibitory pathways is not yet fully understood. One of such inhibitory nodes of the brain is the striatum.

The dorsal striatum is a subcortical part of the forebrain and an input site of the basal ganglia. It coordinates multiple aspects of cognition, including initiation of spontaneous movements, goal-directed and procedural behavior, motivation and reward perception (Balleine & O’Doherty, 2010; Cui et al., 2013; Floresco, 2015; Gremel & Costa, 2013; Tecuapetla et al., 2016; Varin et al., 2023). Striatal projection neurons (SPNs) are GABAergic, and exert a strong inhibition on projection targets and within the striatal microcircuitry (Nelson & Kreitzer, 2014). Main information flows reach striatum from cortex, thalamus and substantia nigra pars compacta (SNpc); and SPNs pass it along the thalamo-cortical loop and then towards the motor nuclei of the brain stem. Striatal projections form two distinct pathways, one initiating, and another inhibiting movement (direct-pathway SPNs, dSPNs and indirect-pathway SPNs, iSPNs). SPNs are located in striosomal and matrix compartments – two neurochemically distinct regions of striatum (Prager & Plotkin, 2019). Matrix-located SPNs form parts of sensorimotor and associative circuits, whereas striosomes contain SPNs that receive input from limbic cortex and project to the dopamine-containing neurons of SNpc.

During early development, SPNs are hyperexcitable and spontaneously active, generating calcium transients and action potentials (Comhair et al., 2018; Dehorter et al., 2011; Kozorovitskiy et al., 2012). Starting from embryonic day 16 (E16), a fraction of neonatal SPNs display spontaneous calcium transients, accompanied by action potentials. Spontaneously active SPNs are present until postnatal day 10 (P10) (Dehorter et al., 2011), after which they become silent and acquire adult-type electrophysiological profile due to expression of potassium inward rectifier channels (KIRs). At the behavioral level, this corresponds to the time when pups start to move actively and exit the nest. Spontaneous activity, however, was detected only in 30% of the neonatal SPNs recorded (Comhair et al., 2018; Dehorter et al., 2011). Neonatal activity of SPNs has also been shown by multi-channel recording *in vivo* (Klavinskis-Whiting et al., 2023). This neonatal activity is organized in bursts, which have different frequency structure, than the adult-type ones, and spreads along the cortico-striato-thalamic loop, probably mediating distinct directional interactions among the basal ganglia structures.

Neuronal activity and subsequent glutamate release are essential for stabilizing cortico-striatal synapses during early postnatal development *in vitro* (Kuhlmann et al., 2021) and *in vivo* (Kozorovitskiy et al., 2012, 2015). Arresting the GABAergic transmission or hyperpolarization of the SPNs by chemogenetic approach during first three weeks resulted in the changes of cortical input in P15 animals, assuming the neonatal activity-dependent modulation of the whole basal ganglia circuit (Kozorovitskiy et al., 2012). However, nothing is known about the activity-dependent maturation of the long-range GABAergic projections.

Taken together, these data show that, like other brain structures, striatum generates and conducts neonatal activity waves, which have frequency characteristics, distinct from adult-type activity. These neonatal waves propagate along the basal ganglia network and support the development and stabilization of synaptic connections. However, two important questions were left open: do all types of SPNs display spontaneous activity in a similar way, and how does neonatal activity guide the development of striatal GABAergic projections – does the “fire together – wire together” rule (Katz & Shatz, 1996) still apply there? We approached these questions here by characterizing the spontaneous activity of neonatal dSPNs and iSPNs, located in either striosomes or matrix; and evaluating the impact of chemogenetic silencing of striosomal SPNs on the development of striato-nigral projections.

## Results

It has been shown before that about 30% of SPNs in the dorsal striatum of one-week old mice fire spontaneous action potentials (Comhair et al., 2018; Dehorter et al., 2011). This low number of spontaneously active SPNs may reflect the heterogeneity in the timing and properties of development of different functional domains of striatum. To check this hypothesis, we evaluated the spontaneous activity and intrinsic excitability of developing SPNs of direct (dSPNs) and indirect (iSPNs) pathways, located either in striosomal or matrix compartments (Fig.1A). dSPNs and iSPNs were recorded in the dorsal striatum of D1-Cre/Ai14 and A2a-Cre/Ai14 animals. Striosomal compartments were recognized in the slices by the higher density of tdTomato signal. Striosomal SPNs are generated two days earlier than matrix SPNs, and consequently, start to express the identity marker proteins earlier (Passante et al., 2008). We confirmed that clusters of dense tdTomato fluorescence in the neonatal striatum correspond to striosomes by immunostaining of striatal slices of one-week-old D1-Cre/Ai14 and A2a-Cre/Ai14 animals for the specific striosomal marker µ-opioid receptor (MOR, Fig. 1B). Specific tdTomato clustering was not visible in D1-Cre/Ai14 slices after P10 and in A2a-Cre/Ai14 slices after P14.

**Figure 1.**
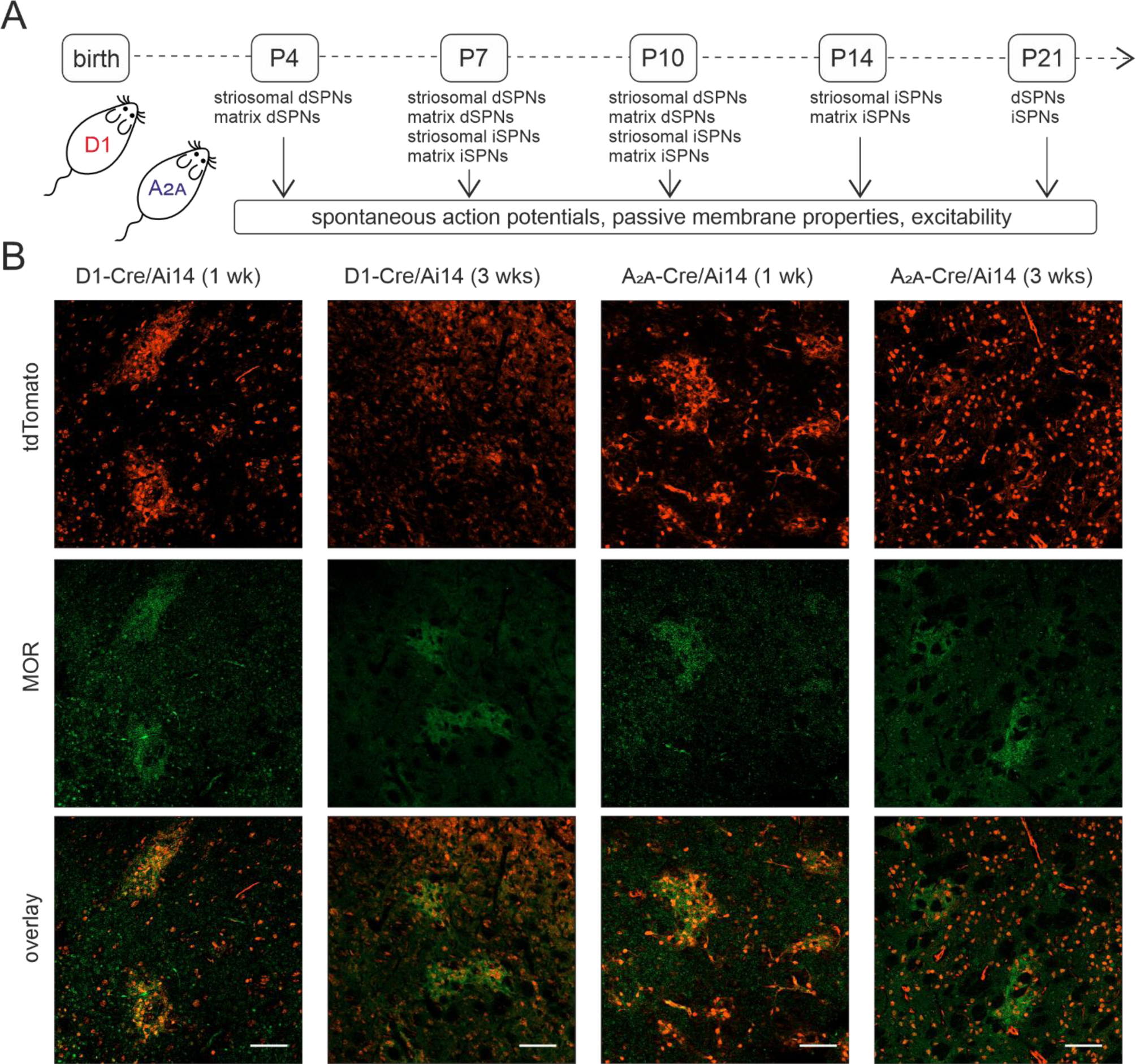
The overview of the experimental approach to describe the activity patterns across striatal functional domains. A. Schematic illustration of the experimental timeline, related to Figures 1-4. D1-Cre/Ai14 and A_2A_-Cre/Ai14 mice, expressing reporter proteins, were used at different ages for the characterization of the spontaneous activity, intrinsic excitability and passive membrane properties. B. Immunostaining of dorsal striatum of D1-Cre and A_2A_-Cre mice of different ages. Striosomes are visible during early postnatal development by denser expression of the tdTomato after Cre recombination under control of Drd1 and A_2A_ promoters. Red: tdTomato; green: µ-opioid receptor staining. Scale bar 100 µm.

Some developing SPNs fire irregular spontaneous action potentials (APs, Fig. 2A). This activity pattern with a stable baseline membrane potential and clear APs, crossing the threshold of 0 mV, was detected in P3-4 dSPNs and P6-7 iSPNs at the earliest. At P4, cells expressing tdTomato after A2a-Cre recombination were present in the striatal slices, but electrophysiological recordings were not stable and no APs, crossing 0 mV were detected. The number of spontaneously active cells decreased with age, striosomal cells being spontaneously active at later developmental stages, compared to matrix ones (Fig. 2B). However, the frequency of APs did not differ between neonatal striosomal and matrix SPNs (dSPNs: striosomes 1.67±0.45 Hz, matrix 2.75±0.71 Hz; p=0.22; iSPNs: striosomes 2.88±0.73 Hz, matrix 1.62±0.57 Hz; p=0.25). Although the cortical projections are present in the tilted slice preparations, we did not see the decrease in generation of spontaneous APs after inhibition of glutamatergic transmission. The frequency of APs recorded in cell-attached configuration from striosomal dSPNs did not change after application of AMPA receptor blocker NBQX (Fig. S1B, baseline 2.25±0.73 Hz; NBQX 2.30±0.68 Hz, p=0.94), and the frequency of APs in iSPNs increased (Fig. S1C, baseline 1.88±0.51 Hz; NBQX 3.60±2.24 Hz, p<0.05). We confirmed the spontaneous activity of developing SPNs in the form of intracellular Ca^2+^ oscillations (Fig 2C for dSPNs and Fig. 2D for iSPNs). The percentage of spontaneously active dSPN and iSPN cells from all active cells in the presence of 8 mM KCl was significantly higher in the striosomal compartment, compared to the matrix one (dSPNs: striosomes 35.00±4.13 %, matrix 21.91±3.13 %, p<0.05; iSPNs: striosomes 45.45±4.82 %, matrix 26.52±2.85 %, p<0.01).

**Figure 2.**
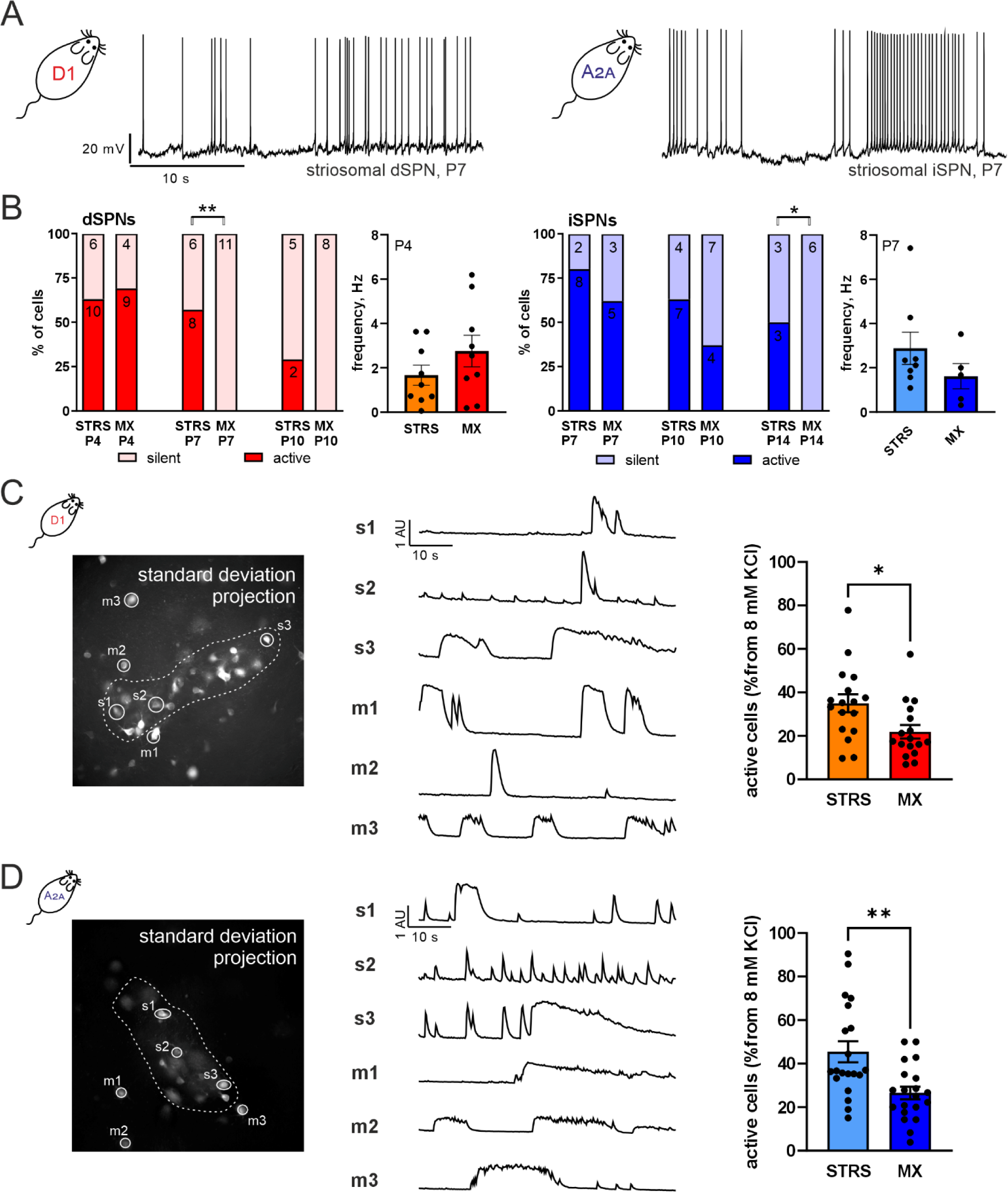
Spontaneous activity in striosomal and matrix dSPNs and iSPNs. A. Example traces of spontaneous APs, recorded by whole-cell patch-clamp from striosomal dSPN of P4 D1-Cre/Ai14 animal and iSPN of P7 A_2a_-Cre/Ai14 animal. B. Percent’s of spontaneously active (with AP firing) and silent (without AP firing) dSPNs and iSPNs in striosomes and matrix of P4-P14 animals. *p<0.05, **p<0.01, Chi-square test. Number of cells is indicated on the bars. Numbers of animals used are: dSPN P4 N=6, P7 N=9, P10 N=4; iSPN P7 N=4, P10 N=5, P14 N=3. Frequency of spontaneous APs in striosomal and matrix dSPNs of P4 animals and striosomal and matrix iSPNs of P7 animals from the same data set as the previous graph. C,D Spontaneous Ca^2+^ oscillations in striosomal and matrix dSPNs (C) and iSPNs (D), recorded in striatal slices of P4 D1-Cre/Ai95D and P7 A_2a_-Cre/Ai95D animals. Striosomal compartment border was marked on the maximal intensity projection of the movie, and ROIs corresponding to the active cells were selected on the standard deviation projection of the movie. Example traces show spontaneous Ca^2+^ oscillations in striosomal and matrix cells of D1-Cre (C) and A_2a_-Cre (D) slices. Percent’s of spontaneously active cells in striosomes and matrix from active cells in 8 KCl conditions are presented as single values per slice and mean±SEM. dSPNs: 17 slices for striosomes, 17 slices for matrix, 3 animals. iSPNs: 20 slices for striosomes, 20 slices for matrix, 5 animals. *p<0.05, **p<0.01, ***p<0.001, unpaired t-test.

The presence of neonatal spontaneous activity, which disappears in adulthood, is usually a feature of developing neurons. At the early developmental stages striosomal cells are a bit more mature than adult cells (Passante et al., 2008). To investigate why the number of spontaneously active SPNs is higher in striosomes, we compared passive membrane properties and intrinsic excitability of striosomal and matrix dSPNs and iSPNs at three age points. It is known that dSPNs are in general less excitable than iSPNs (Gerfen & Surmeier, 2011; Gertler et al., 2008), so in our analysis we focused only on the age- and domain-related effects. Like in the spontaneous activity recordings, we compared dSPN properties at P4, P7 and P10; and iSPN properties at P7, P10 and P14. We also recorded passive and active membrane properties of dSPNs and iSPNs at P21.

With our approach, it was not possible to distinguish striosomal and matrix domains at this late age point, so these data were not included into the statistical analysis. However, data are presented in the tables for the reference (Tables 1 and 3; Fig. 3 and 4). Since in adulthood striosomes occupy roughly 15 % of the total striatal volume (Johnston et al., 1990a), we estimate that our data collected at P21 mostly represent matrix-located neurons.

**Figure 3.**
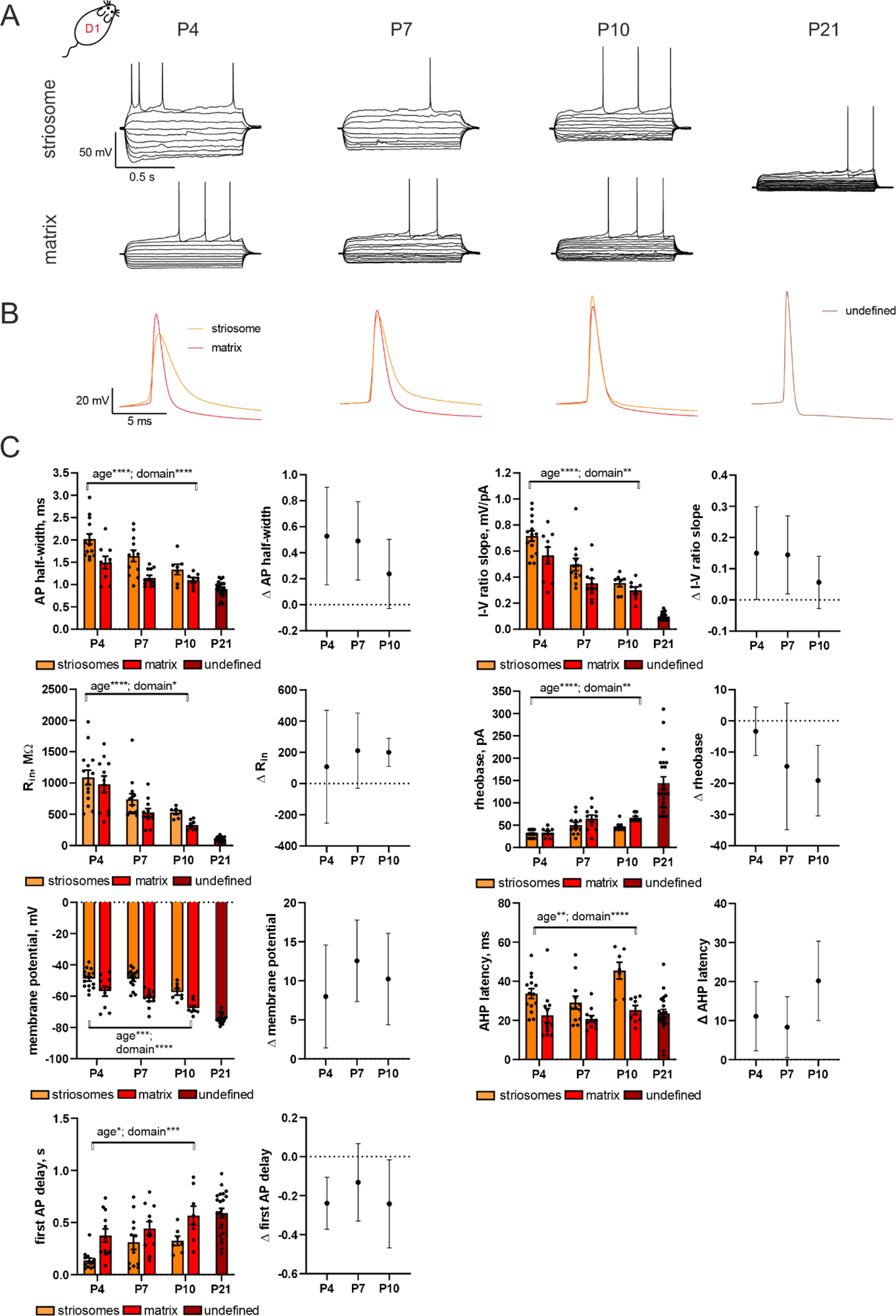
Intrinsic excitability properties of dSPNs during the first three weeks of life. A. Example traces of the membrane potentials after injection of current steps with 10 pA increments of striosomal and matrix dSPNs of dorsal striatum, recorded from P4-P10 D1-Cre/Ai14 animals. Action potentials of P21 dSPN are presented for the comparison (it was not possible to detect the compartment location of the recorded neuron at this age point). B. Example traces of APs at the expanded scale, striosomal and matrix APs are overlayed. C. Membrane properties and AP characteristics, which were significantly different for striosomal and matrix dSPNs. All measured parameters and sample size are presented in Tables S1-2. Data are presented as individual values, mean±SEM, and mean of the difference with 95% CI. *p<0.05, **p<0.01, ***p<0.001, ****p<0.0001, two-way ANOVA.

**Figure 4.**
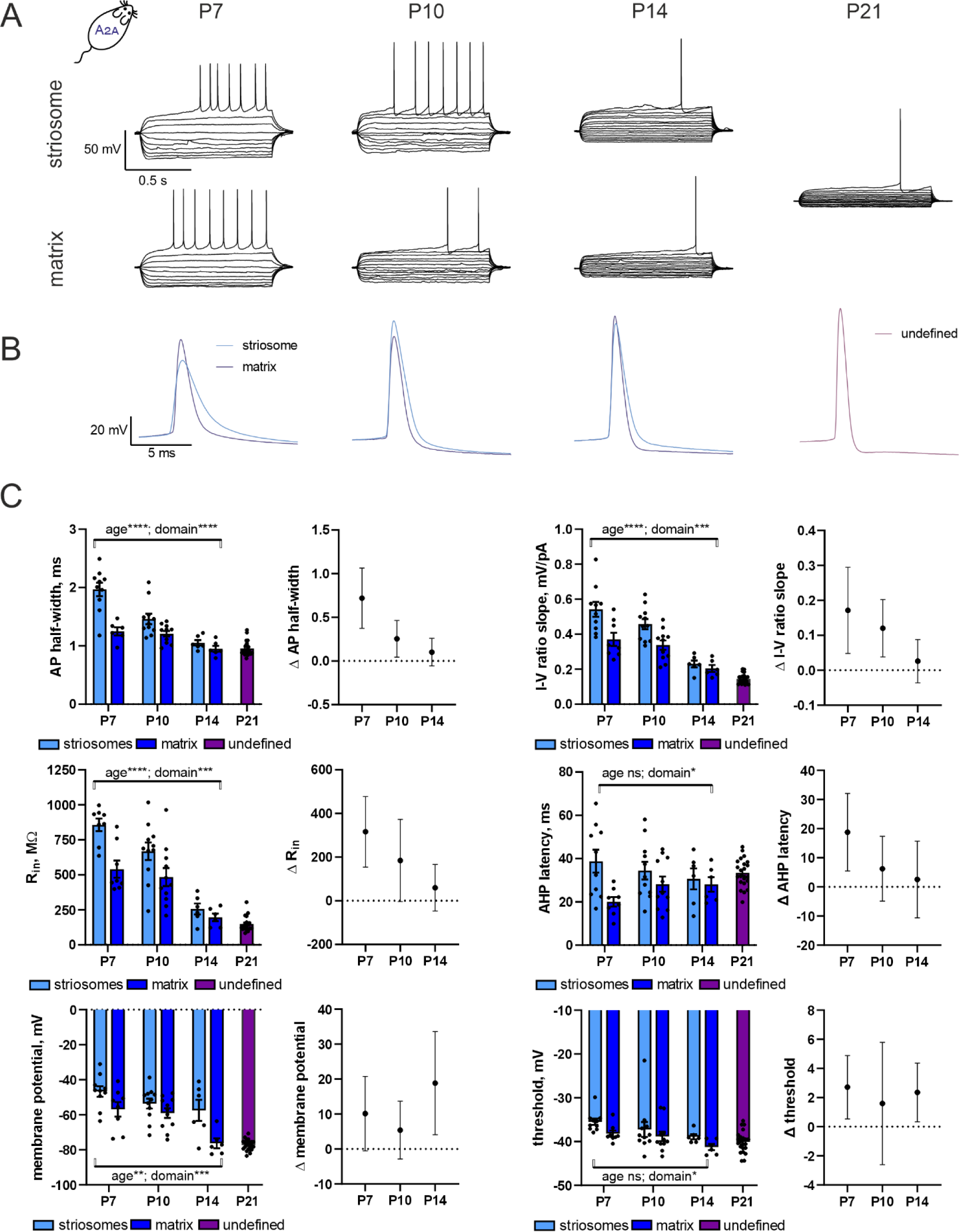
Intrinsic excitability properties of iSPNs during the first three weeks of life. A. Example traces of the membrane potentials after injection of current steps with 10 pA increments of striosomal and matrix iSPNs of dorsal striatum, recorded from P7-P14 A_2A_-Cre/Ai14 animals. Action potentials of P21 iSPN are presented for the comparison (it was not possible to detect the compartment location of the recorded neuron at this age point). B. Example traces of APs at the expanded scale, striosomal and matrix APs are overlayed. C. Membrane properties and AP characteristics, which were significantly different for striosomal and matrix dSPNs. All measured parameters and sample size are presented in Tables S3-4. Data are presented as individual values, mean±SEM, and mean of the difference with 95% CI. *p<0.05, **p<0.01, ***p<0.001, ****p<0.0001, two-way ANOVA.

**Table 1.** Descriptive statistics of the membrane properties of dSPNs.

|  | P4 |  |  | P7 |  |  | P10 |  |  | P21 |
| --- | --- | --- | --- | --- | --- | --- | --- | --- | --- | --- |
|  | striosome | matrix | delta<br>[95%<br>CI] | striosome | matrix | delta<br>[95%<br>CI] | striosome | matrix | delta<br>[95%<br>CI] |  |
| Rin,<br>MΩ | 1088±116.2 | 980.3±130.6 | 108.14<br>[469.99<br>to -<br>253.71] | 739.6±91.5 | 528.1±64.65 | 211.46<br>[452.17,<br>-29.24] | 527.6±31.29 | 327.2±27.57 | 200.38<br>[290.10,<br>110.66] | 102.2±7.14 |
| Cm, pF | 50.56±8.12 | 58.84±8.11 | -8.28 [-<br>31.95,<br>15.39] | 64.66±7.71 | 64.92±6.49 | -0.27 [-<br>21.62,<br>21.08] | 63.74±7.93 | 71.27±6.22 | -7.53 [-<br>29.49,<br>14.43] | 133.50±8.40 |
| Vm, mV | -<br>48.82±1.56 | -56.82±3.15 | 8.0<br>[14.60,<br>1.40] | -<br>48.89±1.69 | -61.44±1.80 | 12.56<br>[17.77,<br>7.34] | -57.36±2.04 | -67.6±1.75 | 10.24<br>[16.10,<br>4.38] | -74.35±0.61 |
| Rheobase,<br>pA | 30.0±2.10 | 33.33±3.33 | -3.33<br>[4.42, -<br>11.09] | 50.5±6.03 | 64.55±7.79 | -14.55<br>[5.74, -<br>34.84] | 47.14±4.21 | 66.25±3.24 | -19.11<br>[-7.80, -<br>30.42] | 144.1±14.17 |
| Threshold,<br>mV | -<br>31.12±0.64 | -36.46±1.02 | 5.34<br>[2.97,<br>7.71] | -<br>33.00±1.44 | -31.58±1.28 | -1.42 [-<br>5.46,<br>2.62] | -33.76±1.63 | -35.09±0.54 | 1.33 [-<br>2.18,<br>4.84] | -37.91±0.85 |
| First AP<br>delay, s | 0.137±0.02 | 0.376±0.06 | -0.239<br>[-<br>0.106, -<br>0.372] | 0.311±0.07 | 0.443±0.07 | -0.132<br>[0.067,<br>-0.331] | 0.325±0.05 | 0.567±0.09 | -0.242<br>[-0.016,<br>-0.468] | 0.591±0.05 |
| AP half-width, ms | 2.02±0.11 | 1.49±0.14 | 0.53<br>[0.9,<br>0.15] | 1.64±0.13 | 1.15±0.06 | 0.49<br>[0.79,<br>0.19] | 1.34±0.12 | 1.10±0.05 | 0.24<br>[0.50, -<br>0.03] | 0.90±0.04 |
| AP amplitude, mV | 60.38±1.43 | 65.89±2.12 | -5.50 [-10.62, -0.39] | 59.28±2.60 | 63.56±2.60 | -4.28 [-11.92, 3.359] | 71.27±3.82 | 68.08±3.81 | 3.20 [-8.497, 14.89] | 75.67±2.51 |
| AHP amplitude, mV | 13.61±1.20 | 12.00±0.92 | 1.61 [-1.597, 4.816] | 10.68±0.75 | 13.55±0.73 | -2.87 [-5.055, -0.6784] | 13.80±0.72 | 12.47±0.52 | 1.33 [-0.5474, 3.209] | 12.95±0.50 |
| AHP latency, ms | 33.69±2.53 | 22.56±3.57 | 11.13 [2.293, 19.97] | 29.09±3.26 | 20.79±1.62 | 8.31 [0.5089, 16.10] | 45.45±4.28 | 25.27±2.34 | 20.18 [10.02, 30.34] | 23.56±2.11 |
| AP frequency slope, Hz/pA | 0.44±0.03 | 0.46±0.03 | -0.019 [-0.118, 0.080] | 0.39±0.04 | 0.30±0.04 | 0.088 [-0.040, 0.215] | 0.42±0.03 | 0.35±0.03 | 0.070 [-0.018, 0.159] | 0.29±0.01 |
| I-V ratio slope, mV/pA | 0.72±0.04 | 0.57±0.06 | 0.15 [0.30, 0.001] | 0.50±0.05 | 0.35±0.04 | 0.14 [0.27, 0.02] | 0.35±0.03 | 0.30±0.03 | 0.06 [0.14, -0.03] | 0.10±0.01 |
|  | striosomal cells | matrix cells | N of animals | striosomal cells | matrix cells | N of animals | striosomal cells | matrix cells | N of animals | n of cells / N of animals |
| sample size | 15 | 10 | 6 | 14 | 10 | 9 | 7 | 7 | 4 | 22/5 |

**Table 2.** Two-way ANOVA of the dSPN membrane properties (P4-P10)

|  | age |  | domain |  | interaction |  |
| --- | --- | --- | --- | --- | --- | --- |
| R<sub>in</sub>, MOhm | F (2, 58) = 18.11 | **P<0.0001** | F (1, 58) = 4.240 | **P=0.0440** | F (2, 58) = 0.1746 | P=0.8402 |
| Cm, pF | F (2, 59) = 1.462 | P=0.2401 | F (1, 59) = 0.6266 | P=0.4318 | F (2, 59) = 0.1662 | P=0.8473 |
| Vm, mV | F (2, 57) = 9.582 | **P=0.0003** | F (1, 57) = 34.42 | **P<0.0001** | F (2, 57) = 0.6950 | P=0.5033 |
| Rheobase, pA | F (2, 55) = 17.08 | **P<0.0001** | F (1, 55) = 8.363 | **P=0.0055** | F (2, 55) = 1.212 | P=0.3056 |
| Threshold, mV | F (2, 55) = 1.773 | P=0.1795 | F (1, 55) = 3.318 | P=0.0740 | F (2, 55) = 4.847 | P=0.0115 |
| First AP delay, s | F (2, 57) = 4.795 | **P=0.0119** | F (1, 57) = 15.96 | **P=0.0002** | F (2, 57) = 0.5485 | P=0.5808 |
| AP half-width, ms | F (2, 55) = 11.15 | **P<0.0001** | F (1, 55) = 19.78 | **P<0.0001** | F (2, 55) = 0.8238 | P=0.4441 |
| AP amplitude, mV | F (2, 55) = 4.771 | **P=0.0123** | F (1, 55) = 1.021 | P=0.3167 | F (2, 55) = 1.372 | P=0.2621 |
| AHP amplitude, mV | F (2, 58) = 0.5641 | P=0.5719 | F (1, 58) = 0.0009651 | P=0.9753 | F (2, 58) = 3.752 | P=0.0293 |
| AHP latency, ms | F (2, 58) = 5.149 | **P=0.0087** | F (1, 58) = 27.18 | **P<0.0001** | F (2, 58) = 1.716 | P=0.1888 |
| AP frequency slope, Hz/pA | F (2, 55) = 4.486 | **P=0.0157** | F (1, 55) = 2.144 | P=0.1488 | F (2, 55) = 1.222 | P=0.3025 |
| I-V ratio<br>slope,<br>mV/pA | F (2, 55)<br>= 24.84 | **P<0.0001** | F (1, 55) =<br>9.849 | **P=0.0027** | F (2, 55)<br>= 0.5782 | P=0.5643 |

**Table 3.**
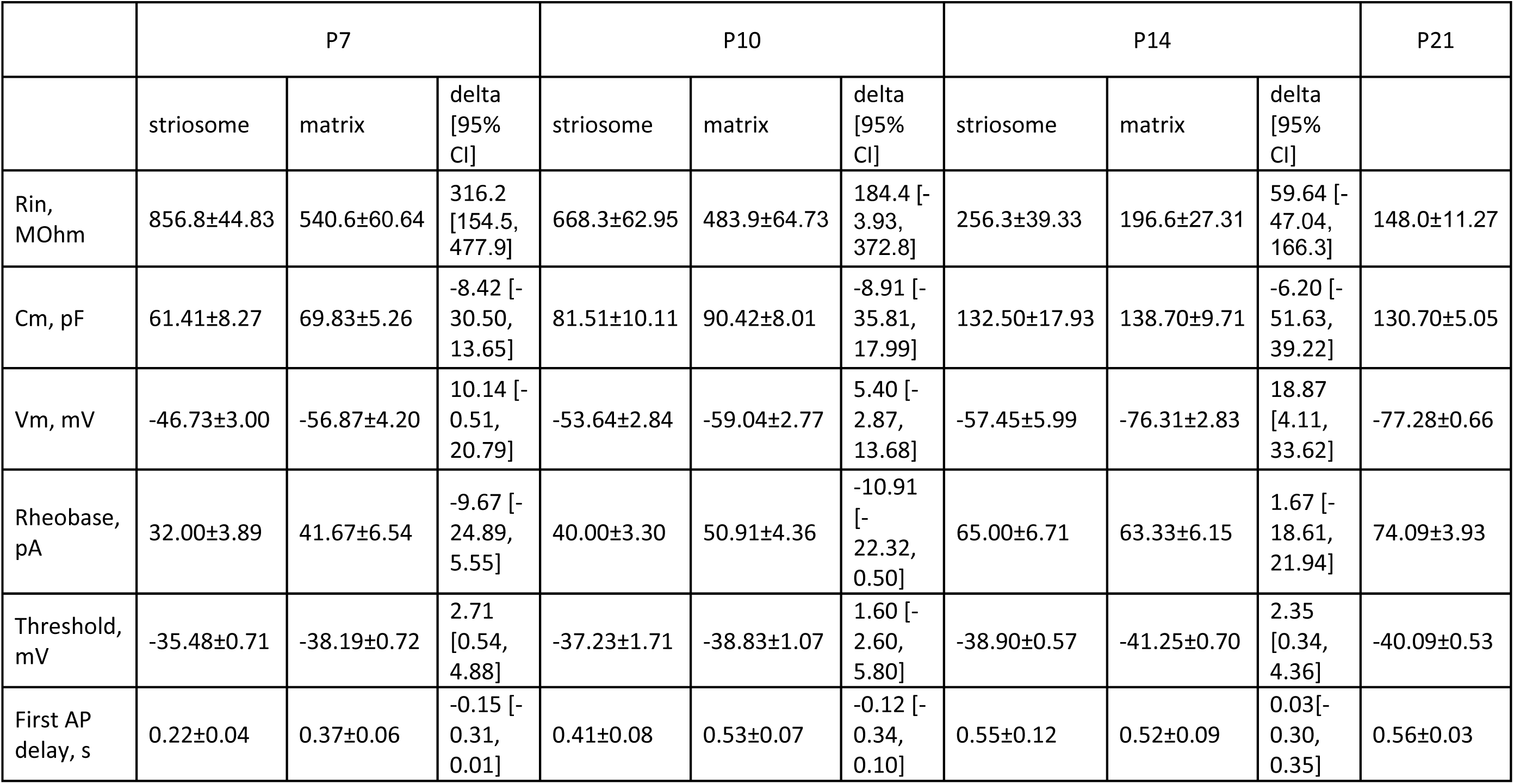
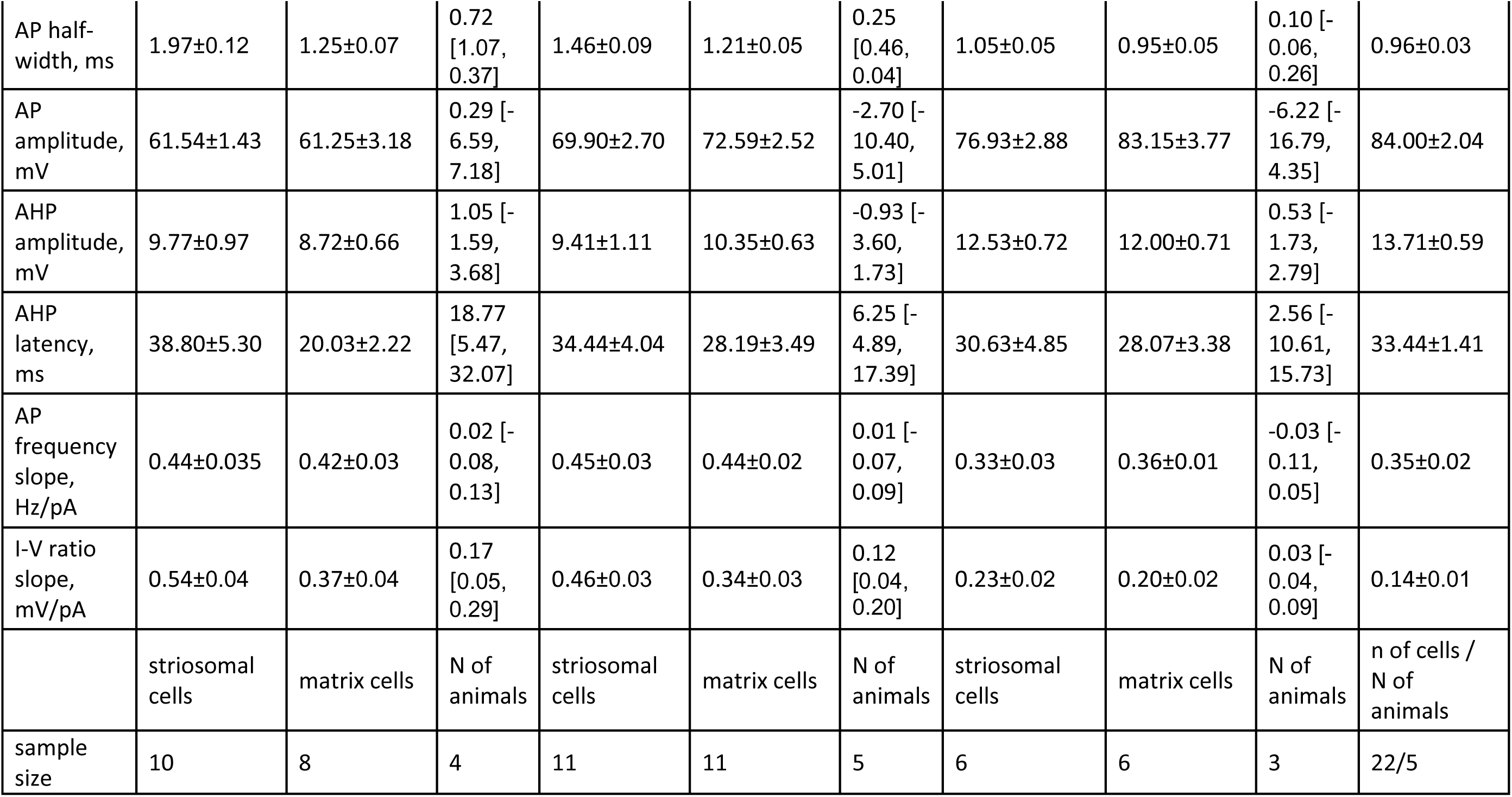
Descriptive statistics of the membrane properties of iSPNs.

|  | P7 |  |  | P10 |  |  | P14 |  |  | P21 |
| --- | --- | --- | --- | --- | --- | --- | --- | --- | --- | --- |
|  | striosome | matrix | delta<br>[95%<br>CI] | striosome | matrix | delta<br>[95%<br>CI] | striosome | matrix | delta<br>[95%<br>CI] |  |
| Rin,<br>MΩ | 856.8±44.83 | 540.6±60.64 | 316.2<br>[154.5,<br>477.9] | 668.3±62.95 | 483.9±64.73 | 184.4 [-<br>3.93,<br>372.8] | 256.3±39.33 | 196.6±27.31 | 59.64 [-<br>47.04,<br>166.3] | 148.0±11.27 |
| Cm, pF | 61.41±8.27 | 69.83±5.26 | -8.42 [-<br>30.50,<br>13.65] | 81.51±10.11 | 90.42±8.01 | -8.91 [-<br>35.81,<br>17.99] | 132.50±17.93 | 138.70±9.71 | -6.20 [-<br>51.63,<br>39.22] | 130.70±5.05 |
| Vm, mV | -46.73±3.00 | -56.87±4.20 | 10.14 [-<br>0.51,<br>20.79] | -53.64±2.84 | -59.04±2.77 | 5.40 [-<br>2.87,<br>13.68] | -57.45±5.99 | -76.31±2.83 | 18.87<br>[4.11,<br>33.62] | -77.28±0.66 |
| Rheobase,<br>pA | 32.00±3.89 | 41.67±6.54 | -9.67 [-<br>24.89,<br>5.55] | 40.00±3.30 | 50.91±4.36 | -10.91<br>[-<br>22.32,<br>0.50] | 65.00±6.71 | 63.33±6.15 | 1.67 [-<br>18.61,<br>21.94] | 74.09±3.93 |
| Threshold,<br>mV | -35.48±0.71 | -38.19±0.72 | 2.71<br>[0.54,<br>4.88] | -37.23±1.71 | -38.83±1.07 | 1.60 [-<br>2.60,<br>5.80] | -38.90±0.57 | -41.25±0.70 | 2.35<br>[0.34,<br>4.36] | -40.09±0.53 |
| First AP<br>delay, s | 0.22±0.04 | 0.37±0.06 | -0.15 [-<br>0.31,<br>0.01] | 0.41±0.08 | 0.53±0.07 | -0.12 [-<br>0.34,<br>0.10] | 0.55±0.12 | 0.52±0.09 | 0.03[-<br>0.30,<br>0.35] | 0.56±0.03 |
| AP half-width, ms | 1.97±0.12 | 1.25±0.07 | 0.72 [1.07, 0.37] | 1.46±0.09 | 1.21±0.05 | 0.25 [0.46, 0.04] | 1.05±0.05 | 0.95±0.05 | 0.10 [-0.06, 0.26] | 0.96±0.03 |
| AP amplitude, mV | 61.54±1.43 | 61.25±3.18 | 0.29 [-6.59, 7.18] | 69.90±2.70 | 72.59±2.52 | -2.70 [-10.40, 5.01] | 76.93±2.88 | 83.15±3.77 | -6.22 [-16.79, 4.35] | 84.00±2.04 |
| AHP amplitude, mV | 9.77±0.97 | 8.72±0.66 | 1.05 [-1.59, 3.68] | 9.41±1.11 | 10.35±0.63 | -0.93 [-3.60, 1.73] | 12.53±0.72 | 12.00±0.71 | 0.53 [-1.73, 2.79] | 13.71±0.59 |
| AHP latency, ms | 38.80±5.30 | 20.03±2.22 | 18.77 [5.47, 32.07] | 34.44±4.04 | 28.19±3.49 | 6.25 [-4.89, 17.39] | 30.63±4.85 | 28.07±3.38 | 2.56 [-10.61, 15.73] | 33.44±1.41 |
| AP frequency slope, Hz/pA | 0.44±0.035 | 0.42±0.03 | 0.02 [-0.08, 0.13] | 0.45±0.03 | 0.44±0.02 | 0.01 [-0.07, 0.09] | 0.33±0.03 | 0.36±0.01 | -0.03 [-0.11, 0.05] | 0.35±0.02 |
| I-V ratio slope, mV/pA | 0.54±0.04 | 0.37±0.04 | 0.17 [0.05, 0.29] | 0.46±0.03 | 0.34±0.03 | 0.12 [0.04, 0.20] | 0.23±0.02 | 0.20±0.02 | 0.03 [-0.04, 0.09] | 0.14±0.01 |
|  | striosomal cells | matrix cells | N of animals | striosomal cells | matrix cells | N of animals | striosomal cells | matrix cells | N of animals | n of cells / N of animals |
| sample size | 10 | 8 | 4 | 11 | 11 | 5 | 6 | 6 | 3 | 22/5 |

We compared two passive membrane properties from the voltage-clamp recordings (input resistance, R_in_ and membrane capacitance, C_m_), membrane potential (V_m_) and nine parameters of the active membrane properties from the current-clamp recordings. Descriptive statistics, estimation statistics and two-way ANOVA values for all the analyzed parameters are presented in Tables 1,2 (dSPNs) and Tables 3,4 (iSPNs). Most of the parameters analyzed changed with the maturation state of the neurons. In dSPNs, during P4-P10 time interval there were no changes in Cm, AP threshold and AHP (afterhyperpolarization) amplitude. However, by P21 C_m_ significantly increased (P10 matrix dSPNs: 71.27±6.22 pF; P21 dSPNs: 133.50±8.40 pF, p=0.004 by t-test). In iSPNs, only AP threshold and AHP latency did not change with age. Two-way ANOVA analysis has shown that domain factor (location of the cell in striosomal or matrix compartment) significantly contributed to the variation in seven membrane parameters of dSPNs and six membrane parameters of iSPNs. To study the impact of the domain factor during three age points, we performed estimation statistics and calculated the mean difference (striosomal value minus matrix value) for each age point analyzed.

In dSPNs, AP half-width, IV ratio slope, R_in_, rheobase, Vm, AHP latency, and first AP delay showed the significant contribution of the domain factor in two-way ANOVA analysis (Table 2). The difference in the AP half-width and I-V ratio slope was maximal in P4 dSPNs (Fig. 3C, Table 1), decreasing with age and disappearing in P10 dSPNs. Differences in the R_in_ and rheobase, on the contrary, appeared only with age (Fig. 3C, Table 1). V_m_, AHP latency and the first AP delay were different in striosomal and matrix dSPNs in all ages analyzed, AHP latency being higher in striosomal neurons, and the first AP delay in matrix neurons (Fig. 3C, Table 1). Higher R_in_, V_m_, AHP latency, lower rheobase, and first AP delay may signify the higher excitability of striosomal dSPNs at later developmental time points; and wider APs and higher IV slope may be linked to the higher excitability at P4-P7.

In iSPNs, the domain factor contributed to the variance in AP half-width, I-V ratio slope, R_in_, AHP latency, V_m_ and AP threshold (Table 4). The difference in the first four parameters was the highest in P7 animals, and disappeared with age, being non-significant in P14 animals (Fig. 4C, Table 2). The difference in the V_m_, on the contrary, only appeared with age, being maximal at P14 (Fig. 4C, Table 2). AP threshold was significantly higher in striosomal iSPNs in P7 and P14 animals (Fig. 4C, Table 2). Higher AP half-width, IV slope, R_in_, and AHP latency may be linked to the increased excitability of striosomal iSPNs at P7, and higher V_m_ and AP threshold is in line with the higher excitability of these cells at later developmental stages.

**Table 4.** Two-way ANOVA of the iSPN membrane properties (P4-P10)

|  | age |  | domain |  | interaction |  |
| --- | --- | --- | --- | --- | --- | --- |
| Rin, MOhm | F (2, 44) = 27.54 | **P<0.0001** | F (1, 44) = 14.06 | **P=0.0005** | F (2, 44) = 1.958 | P=0.1532 |
| Cm, pF | F (2, 46)<br>= 21.47 | **P<0.0001** | F (1, 46)<br>= 0.8945 | P=0.3492 | F (2, 46)<br>= 0.008889 | P=0.9912 |
| Vm, mV | F (2, 46)<br>= 7.882 | **P=0.0011** | F (1, 46)<br>= 15.20 | **P=0.0003** | F (2, 46)<br>= 1.672 | P=0.1991 |
| Rheobase, pA | F (2, 44)<br>= 13.41 | **P<0.0001** | F (1, 44)<br>= 2.391 | P=0.1292 | F (2, 44)<br>= 0.8718 | P=0.4253 |
| Threshold, mV | F (2, 46)<br>= 3.186 | P=0.0506 | F (1, 46)<br>= 5.054 | **P=0.0294** | F (2, 46)<br>= 0.1345 | P=0.8745 |
| First AP delay, s | F (2, 46)<br>= 4.917 | **P=0.0116** | F (1, 46)<br>= 1.650 | P=0.2054 | F (2, 46)<br>= 0.6391 | P=0.5324 |
| AP half-width, ms | F (2, 44)<br>= 20.63 | **P<0.0001** | F (1, 44)<br>= 24.80 | **P<0.0001** | F (2, 44)<br>= 6.270 | P=0.0040 |
| AP amplitude, mV | F (2, 46)<br>= 20.11 | **P<0.0001** | F (1, 46)<br>= 1.579 | P=0.2152 | F (2, 46)<br>= 0.6006 | P=0.5527 |
| AHP amplitude, mV | F (2, 46)<br>= 4.975 | **P=0.0111** | F (1, 46)<br>= 0.07989 | P=0.7787 | F (2, 46)<br>= 0.7476 | P=0.4792 |
| AHP latency, ms | F (2, 46)<br>= 0.1546 | P=0.8572 | F (1, 46)<br>= 6.766 | **P=0.0125** | F (2, 46)<br>= 1.922 | P=0.1578 |
| AP frequency slope, Hz/pA | F (2, 46)<br>= 4.547 | **P=0.0158** | F (1, 46)<br>= 0.001333 | P=0.9710 | F (2, 46)<br>= 0.2918 | P=0.7483 |
| I-V ratio slope, mV/pA | F (2, 46)<br>= 22.65 | **P<0.0001** | F (1, 46)<br>= 14.42 | **P=0.0004** | F (2, 46)<br>= 2.015 | P=0.1449 |

Given that both dSPNs and iSPNs located in striosomes are most likely being spontaneously active than their martrix counterparts, and both cell types have wider APs and higher IV slope, we hypothesize the presence of a common molecular mechanism, keeping these cells active during development. Developing striosomes receive denser cholinergic and dopaminergic innervation, compared to matrix (Graybiel et al., 1981). dSPNs and iSPNs express distinct types of dopaminergic receptors, but they both express muscarinic receptor M1 (Bernard et al., 1992; Yan et al., 2001). Using Oprm1-Cre/Ai14 animals, we confirmed that choline acetyltransferase (ChAT) immunoreactivity is denser in neuropil of striosomes of P7 animals, compared to matrix neuropil (Fig. 5A, striosomes 59.72±2.48 A.U., matrix 47.89±1.83 A.U., p<0.0001). This ratio changes during development, and in the adult striatum ChAT immunoreactivity is higher in the matrix compartment (striosomes 61.39±3.07 A.U., matrix 64.30±3.16 A.U., p<0.0001). Application of the M1 inhibitor VU0255035 significantly reduces the frequency of spontaneous APs in striosomal iSPNs (Fig. 5C, baseline 2.53±0.29 Hz, VU0255035 1.69±0.31 Hz, p<0.05), but not in matrix ones (baseline 1.92±0.54 Hz, VU0255035 2.19±0.57 Hz, p=0.70). Application of VU0255035 did not change the frequency of spontaneous APs both in striosomal dSPNs (baseline 1.60±0.36 Hz, VU0255035 1.36±0.46 Hz, p=0.62) and in matrix dSPNs (baseline 1.76±0.54 Hz, VU0255035 1.84±0.47 Hz, p=0.80). Inhibition of M1 receptors affected the occurrence of Ca^2+^ spikes in striosomal, but not matrix iSPNs and dSPNs (Fig. 5D). The number of Ca^2+^ spikes was significantly decreased in striosomal iSPNs (baseline 73.28±7.89 events/min, VU0255035 30.97±4.81 events/min, p<0.0001), but not affected in matrix ones (baseline 41.77±6.71 events/min, VU0255035 37.35±6.14 events/min, p=0.30). We also observed a decrease in the number of Ca^2+^ spikes in striosomal dSPNs (baseline 50.17±8.35 events/min, VU0255035 35.72±9.56 events/min, p<0.001), although the size of the effect was smaller. VU0255035 did not affect Ca^2+^ spikes in matrix dSPNs (baseline 58.18±8.23 events/min, VU0255035 48.29±10.15 events/min, p=0.07).

**Figure 5.**
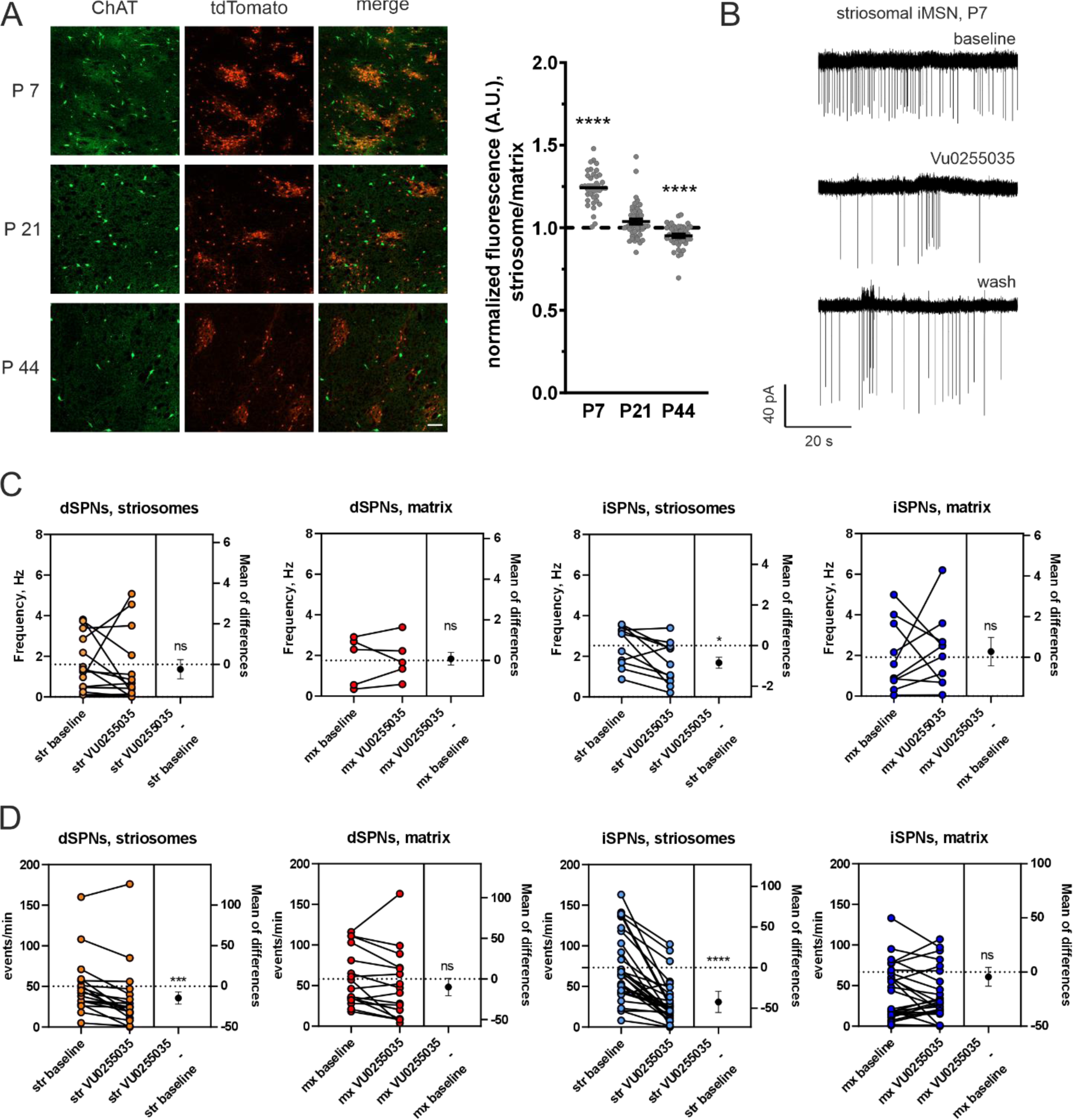
Cholinergic modulation of the neonatal spontaneous activity. A. Immunostaining for ChAT of the dorsal striatal slices of Oprm1-Cre/Ai14 animals at the age of P7, P21 and P44. Green: ChAT staining, Red: tdTomato. Scale bar 100 µm. The density of fluorescent signal in striosomal domains was normalized to the density of fluorescent signal in the matrix domain of the same image. Data are presented as individual values and mean±SEM; P7: 37 images/5 animals; P21: 49 images/7 animals; P44: 44 images from 3 animals. Statistical analysis: Non-normalized fluorescence density of green signal in striosomal regions was compared to matrix ones, ****p<0.0001, Wilcoxon test. B. Example traces of APs, recorded in the cell-attached mode from the striosomal iSPN of the P7 A_2A_-Cre/Ai14 animal during baseline, application of the inhibitor of muscarinic M1 receptor VU0255035 (5 µM) and wash-out. C. Quantification of AP frequency before and during application of VU0255035 in P4-5 striosomal and matrix dSPNs and P6-7 striosomal and matrix iSPNs. Sample size, dSPNs: 14 striosomal cells, 5 matrix cells, 5 animals; iSPNs: 11 striosomal cells, 10 matrix cells, 8 animals. Data are presented as individual values and mean of the difference with 95% CI, *p<0.05, paired t-test. D. Quantification of the number of Ca^2+^ spikes per minute before and during application of VU0255035 in P4-5 striosomal and matrix dSPNs and P6-7 striosomal and matrix iSPNs. Sample size, dSPNs: 18 slices, 3 animals; iSPNs: 29 slices, 6 animals. Data are presented as individual values and mean of the difference with 95% CI, ***p<0.001, ****p<0.0001, paired t-test.

It is generally thought that spontaneous activity of developing neurons is needed for stabilization of synaptic contacts. We tested if spontaneous activity of SPNs affects the maturation of their long-range projections. Since spontaneous activity is more profound in the striosomal compartment, we decided to study the activity-dependent development of the direct projection, originating from striosomes and ending on the dopaminergic (DA) neurons of SNpc, called striosome-dendron bouquets (Crittenden et al., 2016; McGregor et al., 2019). We inhibited striosomal SPNs during the second week of life using a chemogenetic approach and evaluated the striato-nigral connectivity in adult animals (Fig. 6A). Striosomal SPNs of Oprm1-Cre animals and wild-type littermates were transfected with pAAV-hSyn-DIO-HA-hM4D(Gi)-IRES-mCitrine at the age of P1. At P10, mCitrine was densely expressed in the striosomal areas of Oprm1-Cre animals (Fig. S2B), and bath application of CNO blocked spontaneous APs (Fig. S2C, activity block verified using 3 spontaneously active cells). After transfection, animals were chronically injected with CNO or saline during P6-P14, and the GABAergic functional innervation of DA neurons was evaluated in adult animals (P52-79). DA neurons of SNpc were located by their position in the slice, large cell body and the presence of I_h_ current, and the identity of recorded and biocytin-filled neurons was confirmed by *post-hoc* immunostaining for tyrosine hydroxylase (TH). The frequency of miniature inhibitory postsynaptic currents (mIPSCs) was significantly reduced in Oprm1-Cre animals which had been chronically injected with CNO (1.82±0.33 Hz), compared to Oprm1-Cre saline control (3.81±0.5 Hz, p<0.05), wild-type saline control (4.11±0.73 Hz, p<0.05) or wild-type CNO control (4.19±0.59 Hz, p<0.05). Injection of CNO did not change the amplitude of mIPSCs of Oprm1-Cre animals (17.71±1.78 pA, Fig. 6B,C), compared to controls (Oprm1-Cre+saline 14.32±1.09 pA, p=0.5; wild-type+saline 15.5±1.53 pA, p=0.83; wild-type+CNO 20.04±2.4 pA, p=0.72). The capacitance of DA neurons was also not affected (Oprm1-Cre+CNO 83.7±7.87 pF; Oprm1-Cre+saline 84.17±4.41 pF, p>0.99; wild-type+saline 80.95±6.3 pF, p=0.98; wild-type+CNO 84.69±7.26 pF, p=0.99). However, chronic injection of CNO significantly elevated the R_in_ of DA cells in Oprm1-Cre animals (877.4±156.1 MOhm), compared to controls (Oprm1-Cre+saline 490.5±69.78 MOhm, p<0.05; wild-type+saline 298.7±23.09 MOhm, p<0.001; wild-type+CNO 473.8±71.13 MOhm, p<0.01). CNO treatment and activity silencing in the neonatal period did not affect the shape of the DA neuron dendritic tree, analyzed by imaging of biocytin-filled DA neurons from all four experimental groups (Fig. S3A, B).

**Figure 6.**
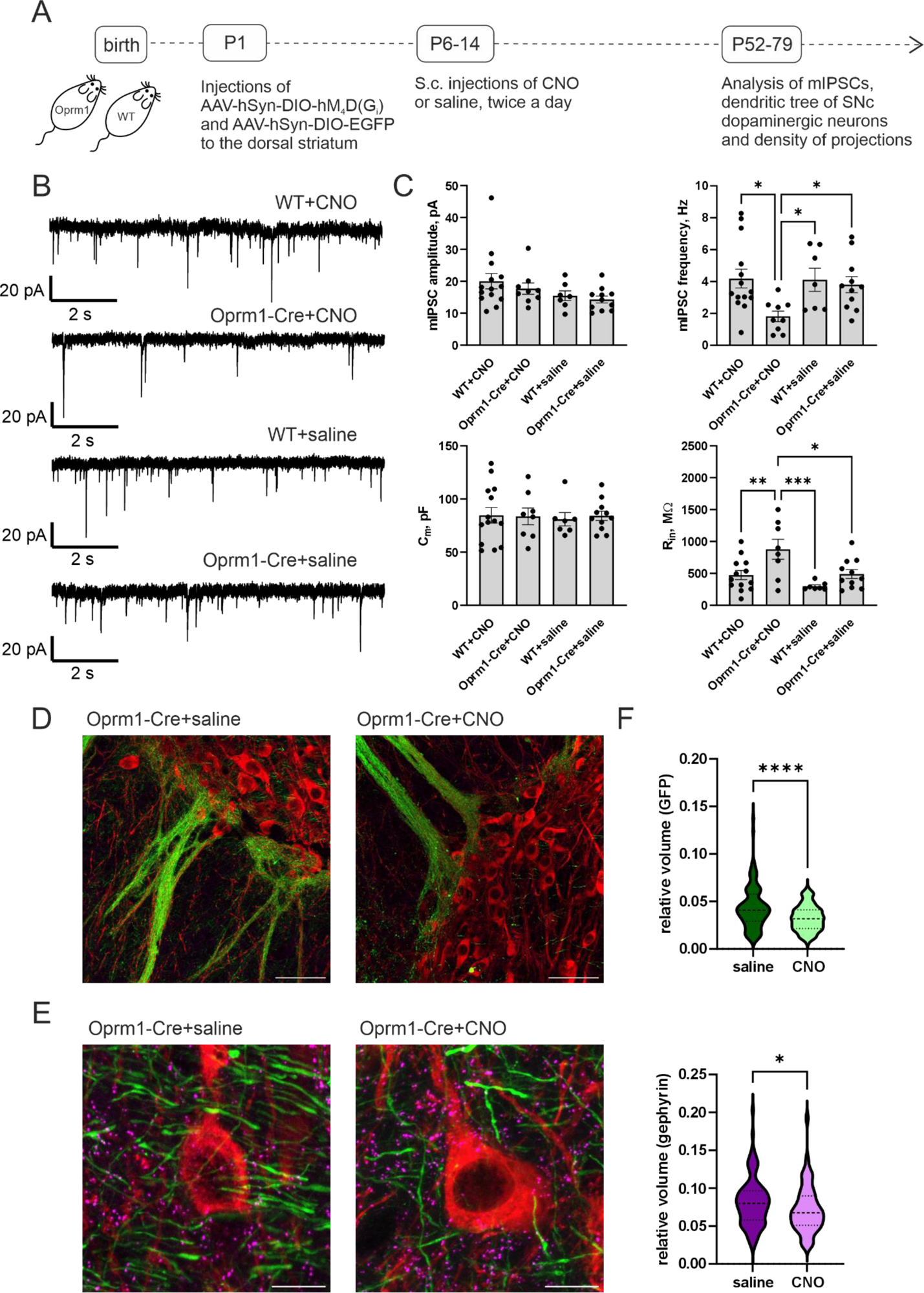
Suppression of the neonatal spontaneous activity during the second week of life causes the decrease in functional GABAergic innervation of SNpc. A. Schematic illustration of the experimental timeline. Oprm1-Cre animals and wild-type littermates received cranial injection of AAV-hSyn-DIO-hM_4_D(G_i_) alone or together with AAV-hSyn-DIO-EGFP into dorsal striatum at P1. At P6-P14 animals received s.c. injections of CNO or saline twice a day. Animals were sacrificed at the age P52-79 for the electrophysiology and imaging experiments. B. Representative traces of mIPSCs recorded from SNpc DA neurons of experimental animals, treated either by CNO or by saline. C. Amplitude and frequency of mIPSCs and capacitance and input resistance of recorded neurons. Sample size, cells/animals: WT CNO 14/9, Oprm1-Cre CNO 9/6, WT saline 7/4, Oprm1-Cre saline 11/7. Data are presented as individual values and mean ± SEM. *p<0.05, ***p<0.001, one-way ANOVA with Holm-Šídák’s multiple comparisons test. D. Example images of striosome-dendrone bouquets of Oprm1-Cre animals after neonatal CNO or saline treatment. Red: TH staining, green: GFP. Scale bar 50 µm. E. Example images of individual DA neurons at higher magnification from the same slices as in D. Red: TH staining; green: GFP; purple: gephyrin staining. Scale bar 10 µm. F. Quantification of the relative GFP and gephyrin volumes from the images of individual DA cells. Sample size, saline: 100 cells/20 slides/5 animals; CNO: 121 cell/24 slides/6 animals. Data are presented as violin plots, median: long-dashed line, quartiles: short-dashed lines. *p<0.05, ****p<0.0001, t-test.

Our electrophysiological data show that chronic suppression of striosomal SPN activity during the second week of life leads to reduced GABAergic drive to DA neurons of the SNpc. To confirm that this effect may be related to reduced innervation of DA neurons by striosomal SPNs, we transfected Oprm1-Cre animals with pAAV-hSyn-DIO-HA-hM4D(Gi)-IRES-mCitrine together with pAAV-hSyn-DIO-EGFP at the age of P1. Transfected animals received chronic CNO or saline injections similarly as for electrophysiological recordings, and animals were sacrificed for immunostaining at the age of P52-79. Coronal slices, containing SNpc, were stained for the DA marker TH and the marker of GABAergic synapses gephyrin. The structure of striosome-dendron bouquets was clearly visible in both animal groups (receiving either CNO or saline injection, Fig. 6D). With our slice preparation, it was difficult to ensure that the whole striosome-dendron bouquet structure was imaged. Therefore, the density of GFP and gephyrin signals were measured around TH-positive cell bodies (Fig. 6 E). GFP signal was measured as voxel volume of green signal per total voxel volume of image, and gephyrin signal – as number of dots per image volume. Injection of CNO caused a significant reduction in the density of GFP signal, which marks striosomal projections (saline 0.044±0.002 RV, CNO 0.032±0.001 RV, p<0.0001). The density of gephyrin staining was also reduced (saline 0.081±0.003 RV, CNO 0.072±0.003 RV, p<0.05), although the effect size was smaller than for GFP.

Our data suggest that reducing the activity of developing SPNs causes a decrease in the density of their projections. To confirm that this effect is specific for the neonatal period, we repeated the activity suppression protocol, but administered CNO to juvenile animals (during P21-29 – the period, where SPNs are already tonically silent, Fig. 7A). Activity suppression during the juvenile period did not decrease, but even increase the frequency of mIPSCs recorded from adult DA neurons (Oprm1-Cre+CNO 5.24±3.93 Hz, wild-type+CNO 3.93±0.35 Hz, p<0.05), and it did not affect the amplitude of these events (Fig. 7B,C; Oprm1-Cre+CNO 26.04±1.57 pA, wild-type+CNO 20.98±1.95 pA, p=0.11). There were no differences in the Rin of DA neurons (Oprm1-Cre+CNO 427.1±80.51 MOhm, wild-type+CNO 403.0±51.43 MOhm, p=0.8), but the C_m_ of DA neurons increased in Oprm1-Cre animals after the juvenile treatment with CNO (Oprm1-Cre+CNO 107.6±9.16 pF, wild-type+CNO 78.92±4.81 pF, p<0.01).

**Figure 7.**
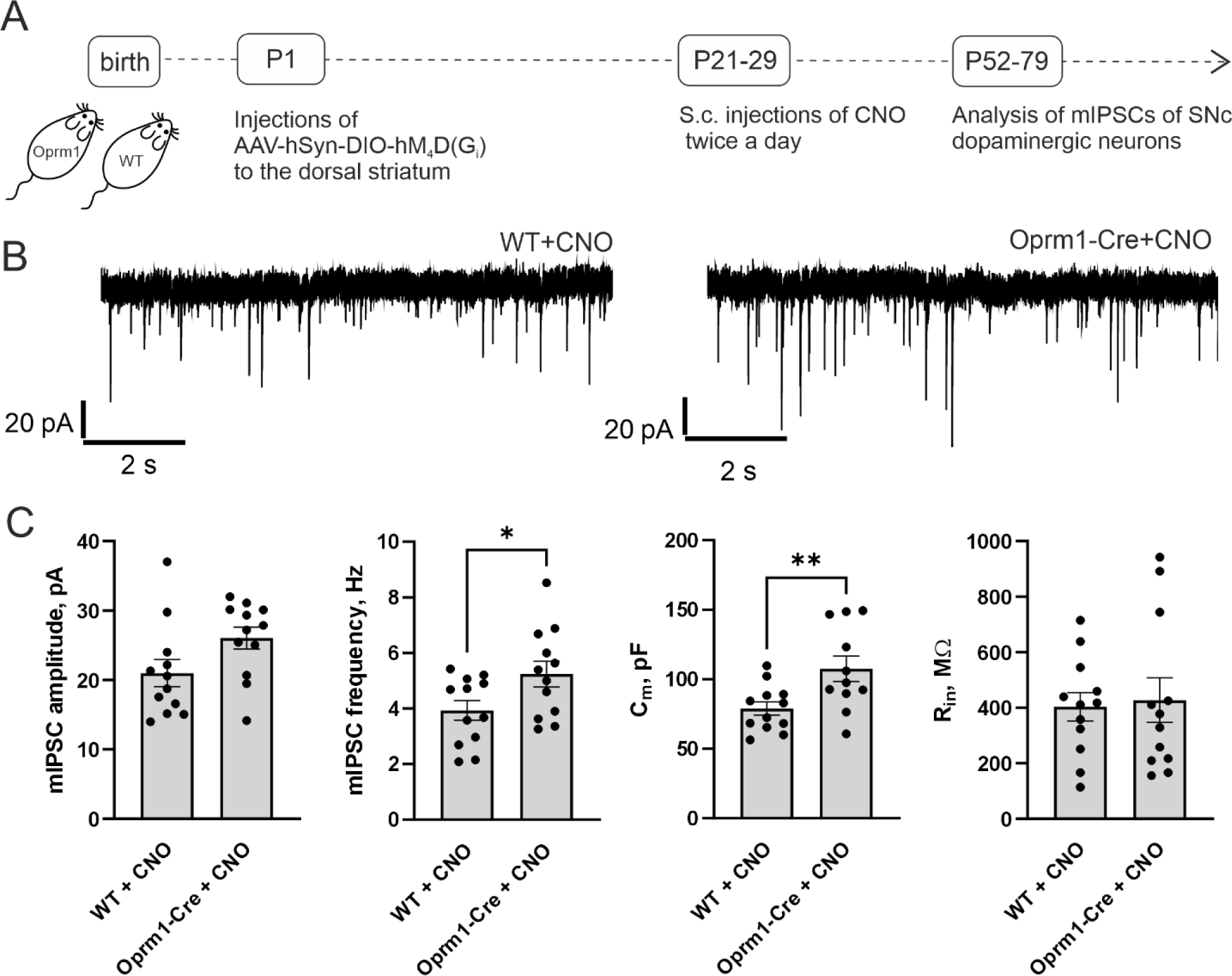
Suppression of striosomal activity in juvenile age does not cause the decrease of the functional GABAergic innervation of SNpc DA neurons. A. Schematic illustration of the experimental timeline. Oprm1-Cre animals and wild-type littermates received cranial injection of AAV-hSyn-DIO-hM_4_D(G_i_) into dorsal striatum at P1. At P21-P29 animals received s.c. injections of CNO twice a day. Animals were sacrificed at the age P52-79 for the electrophysiology experiments. B. Representative traces of mIPSCs recorded from SNpc DA neurons of Oprm1-Cre or wild type animals, treated by CNO. C. Amplitude and frequency of mIPSCs and capacitance and input resistance of recorded neurons. Sample size, cells/animals: WT CNO 12/3, Oprm1-Cre CNO 12/6. Data are presented as individual values and mean ± SEM. *p<0.05, **p<0.01, t-test.

## Discussion

Cortical and thalamic excitatory inputs contribute to the spontaneous firing of developing SPNs (Dehorter et al., 2011; Kozorovitskiy et al., 2015). However, not all neonatal bursts in the striatum are coherent with cortical activity (Klavinskis-Whiting et al., 2023). It is very likely that similar to the multiple origins of neonatal spontaneous activity in other brain regions (Blankenship & Feller, 2010; Luhmann et al., 2016; Molnár et al., 2020), the striatum also generates the activity in the intrinsic networks. Our data show that there are spontaneous calcium oscillations and spontaneous action potentials generated by a fraction of SPNs, an observation that applies to all four types of SPNs we were analyzing (strisosmal dSPNs, strisomal iSPNs, matrix dSPNs, matrix iSPNs). It should be noted that striatum contains a fraction of SPNs, co-expressing D1 receptor and D2 receptor (Bonnavion et al., 2023). D1R/D2R co-expressing cells accounted for 6-7% of all cells, and in our experiments, we will see these cells as dSPNs. With our approach, we cannot distinguish if this activity is purely spontaneous or driven by cortical inputs. We have not seen any decrease in spontaneous AP frequency upon inhibition of AMPA receptors, but it is possible that the cortex is contributing to SPN activity by other means. Regardless, the important finding is that the electrical behavior of neonatal SPNs drastically differs from adult ones, promoting the activity through the basal ganglia circuit.

The general trend is that both dSPNs and iSPNs located in striosomes are more likely to be spontaneously active. This is especially evident when comparing the presence of spontaneous activity in different age groups. Striosomal SPNs stay spontaneously active longer during development, compared to matrix SPNs. Since striosomes occupy roughly 15 % of the striatal volume (Johnston et al., 1990b), higher excitability of these neurons may explain earlier data of SPNs of any type saying that only 30% of neonatal SPNs are spontaneously active.

If we consider neonatal activity as a feature of developing neurons, we could assume that early born neurons (like striosomal SPNs (Passante et al., 2008)) will develop lower membrane potential sooner than matrix SPNs. Since that is not the case, it is likely that there are additional intrinsic excitability mechanisms keeping striosomal SPNs active for a longer time. The development of intrinsic excitability in dSPNs and iSPNs has been addressed in detail before (Gertler et al., 2008; Krajeski et al., 2019; Lieberman et al., 2018). The electrical properties of both populations of the projection neurons develop in parallel during the first three postnatal weeks, and the excitability differences between dSPNs and iSPNs are evident from the early developmental stages. Here, we focus more on the comparison of striosomal vs. matrix SPNs.

Striosomes contain both dSPNs and iSPNs, dSPNs being roughly 70 % of the total SPN count in striosomes of the adult striatum (McGregor et al., 2019; Miyamoto et al., 2018). Our analysis of the membrane properties of striosomal and matrix dSPNs and iSPNs showed that striosomal projection neurons may be in general more excitable, compared to the matrix ones. However, the specific mechanisms may differ, depending on the developmental stage. Striosomal SPNs in the adult striatum are more excitable than matrix SPNs (Crittenden et al., 2017; McGregor et al., 2019; Prager et al., 2020; Smith et al., 2016). These studies, either not distinguishing between the pathway-specific populations, or focusing on dSPNs only, showed that striosomal neurons have higher V_m_ and R_in_ and lower rheobase. In our dataset, we see similar differences in striosomal vs. matrix dSPN properties. Differences in the R_in_ and rheobase appeared by P10, and differences in the V_m_, first AP delay, and AHP latency were evident in all time points analyzed. The excitability properties of adult striosomal and matrix iSPNs have not been studied before. We have shown that P14 striosomal iSPNs have higher V_m_ and AP threshold, which may also contribute to the higher excitability of striosomal SPNs in adulthood (Crittenden et al., 2017).

Although the abovementioned parameters may explain the higher excitability of striosomal SPNs in adulthood, they are most likely not responsible for the higher occurrence of the spontaneously firing AP cells in early developmental stages. During very early time points, both striosomal dSPNs and iSPNs have wider APs and steeper I-V ratio slope, compared to the corresponding matrix cells. In addition, neonatal striosomal iSPNs have higher R_in_ and AHP latency, compared to the matrix iSPNs. It has been shown that steep IV slope of SPNs during development correlates with the delayed expression of inward rectifying potassium channels (KIRs) (Dehorter et al., 2011; Lieberman et al., 2018). The adult-type properties of KIRs appear in SPNs only after the age of two weeks, at the time when pups start to leave the nest and actively explore the surrounding environment (Dehorter et al., 2011). The dynamics of the AP waveform development has not been studied in striatum before. The membrane repolarization after an AP is usually mediated by Kv1, 3 and 4 channels and large conductance calcium-activated potassium channels (Bean, 2007). It is possible that the expression or activity of both KIRs and voltage- or calcium-dependent potassium channels increase in all SPNs during the first weeks of life, but their activity is specifically downregulated in striosomal SPNs, contributing to the higher spontaneous activity in these cells.

Potassium and calcium conductance are strongly regulated by metabotropic receptors, coupled to G proteins. Striatal SPNs receive a dense cholinergic (from local cholinergic interneurons, ChINs) and DA (long-range projections from SNpc) innervation, which regulates their excitability. During development, both DA and cholinergic synapses appear in the striatum during the first postnatal week (Ferrari et al., 2012; Schlösser et al., 1999). During the early development of the cat, both DA and cholinergic innervation has been shown to be denser in striosomes (Graybiel, 1984). Based on these data, both neuromodulators may be responsible for the higher excitability of striosomal SPNs during mammalian development. However, since DA affects the maturation of SPNs only starting from P18 (Lieberman et al., 2018), we decided to focus on the role of muscarinic neuromodulation of striosomal vs matrix SPNs. We confirmed that during early development, cholinergic innervation is denser in striosomal regions, and this is changing with maturation, shifting to denser innervation of matrix regions in the adult brain.

Both dSPNs and iSPNs express muscarinic M1 receptors – metabotropic receptors, coupled to the G_q_ pathway (Bernard et al., 1992; Yan et al., 2001). Initiation of G_q_-coupled signaling results in the activation of PLCβ, degradation of PI(4,5)P_2_, and release of IP_3_. IP_3_ release induces the efflux of Ca^2+^ from endoplasmic reticulum, while a decrease in the PI(4,5)P_2_ membrane content shuts down the function of many potassium channels, including KIRs and BKs (Hille et al., 2015). Activation of M1 receptors elevates the excitability of presumably both types of SPNs by suppressing KCNQ channel current (Shen et al., 2005) and inhibits Kir2 channels only in iSPNs (Shen et al., 2007). We found that pharmacological inhibition of M1 receptors effectively decreases the frequency of spontaneous APs and calcium oscillations in striosomal iSPNs. The effect on dSPNs was much smaller. In those cells, the inhibition of M1 receptors affected only the frequency of spontaneous calcium oscillations, and the effect size was small. It may be so, that some other mechanism contributes to the elevated excitability of striosomal dSPNs, together with M1 receptor.

Finally, we wanted to investigate if spontaneous activity of SPNs is needed for the stabilization of their projections. We approached this task by chemogenetically reducing the probability of SPNs to fire the APs during defined time window in early development. We selected to focus on striosomal SPNs because these neurons generate spontaneous APs for a longer developmental period than matrix ones. Striosomes contain both dSPNs and iSPNs. Similar to matrix iSPNs, striosomal iSPNs project to globus pallidus (Fujiyama et al., 2011). However, striosomal dSPNs are the only striatal cells projecting to DA neurons in SNpc (Fujiyama et al., 2011). These projections cross substantia nigra pars reticulata in a dense bundle and innervate DA neurons in SNpc, forming a morphological structure called “striosome-dendron bouquet” (Crittenden et al., 2016; McGregor et al., 2019). During development, striatonigral projections are formed embryonically (peak at E12), but the critical time window for stabilization of these projections has not been described yet (Matsushima & Graybiel, 2020). Striosomal dSPNs form inhibitory GABAergic synapses onto DA neurons, and activation of striosomal input inhibits spontaneous firing of DA neurons (Evans et al., 2020; McGregor et al., 2019). We demonstrated that chemogenetic silencing of striosomal SPNs in the neonatal period of development caused the weakening of functional GABAergic connectivity to DA cells and decreased the density of striosomal projections to SNpc and gephyrin puncta on DA cells, even though DA neurons also receive GABAergic innervation from substantia nigra pars reticulata (Paladini & Tepper, 2016) and globus pallidus (Evans et al., 2020).

Mechanisms underlying the activity-dependent stabilization of long-range GABAergic projections are still unclear. Most of the previous studies concentrated on the synapses formed by interneurons. During the early development in the cortical structures GABA released by interneurons is depolarizing due to low efficiency of Cl^-^ extrusion from the cell by K-Cl cotransporter KCC2 (Oh & Smith, 2019). In the mechanism stabilizing synapses, the activation of GABA_A_ receptors causes depolarization and Ca^2+^ entrance at the post-synaptic site, and the stabilization of the synapse by cAMP-dependent mechanism of recruiting the scaffolding protein gephyrin to the postsynaptic site. DA neurons do not express KCC2, and Cl^-^ extrusion is due to activation of sodium-dependent anion (Cl^-^-HCO3^-^) exchanger and voltage-sensitive chloride channel 2 (CIC2) (Gulácsi et al., 2003). The Cl^-^ reversal in DA neurons is significantly less negative than in neighboring GABAergic neurons (Gulácsi et al., 2003), but striatal input is still inhibitory in adult SNpc (Evans et al., 2020). The development of Cl^-^ extrusion mechanisms in DA neurons has never been studied, and it is difficult to directly apply the Cl^-^-based mechanisms in activity-dependent stabilization of striosome projections to DA neurons. It was found that the development of striatonigral projections is negatively controlled by cannabinoid receptor 1, although the exact mechanism is unknown (Crittenden et al., 2022). More research is needed to address that in detail.

In contrast to what was seen after neonatal suppression of striosomal SPNs, chemogenetic inhibition of striosomal SPNs during juvenile period resulted in higher mIPSC frequency and C_m_ of DA cells in the adult animals. Thus, the activity-dependent stabilization of striosomal projections has different developmental phases, and the long-term outcome of perturbations in these processes depends on the developmental period when they occur. Suppression of activity later in life may cause some homeostatic changes, directed to maintain the overall activity levels at the silenced synapses (Chen et al., 2022). This is important for studies of neurodevelopmental disorders in animal models when gene expressions or neuronal activity are affected in a time-restricted manner.

## Acknowledgements

Funding for this research was provided by Sigrid Juselius Foundation (T.T. and S.M., Finland), Research Council of Finland (former Academy of Finland; #330298, M.R.; #326045, T.T.; #324548, S.M.), Jane and Aatos Erkko Foundation (S.M., Finland), FRS-FNRS (S.N.S., A.K.E., Belgium) and Action de Recherche Concertée (FWB, S.N.S., A.K.E., Belgium). A.K.E. is a research director of the FRS-FNRS and a WELBIO Investigator. We thank Prof. Konstantinos Meletis and Dr. Anastasia Ludwig for sharing Oprm1-Cre and Ai95D mouse lines with us. Open access funded by Helsinki University Library.

## Author Contributions

B.K., P.S., A.D., M.R. and S.M. designed and conducted the experiments and analyzed the data. S.N.S. and A.K.E. generated transgenic mouse line. T.T. helped with conceptualization of the study and supported the initial stages of the project. S.M. conceptualized the study, S.M. and B.K. wrote the paper.

## Methods

### Animals

All animal experiments were conducted according to the ethical guidelines (in accordance with EU directive 2010/63/EU) provided by the University of Helsinki. The animals were kept under a 12:12 h light/dark cycle (light on at 6AM) and were provided with *ad libitum* access to food and water. Both female and male mice were used for all experiments. The reduction and refinement of the animal experiments was achieved by an *ad hoc* power calculation (N) and designing paired experiments if possible. Mouse strains used in the study are: C57Bl/6JRccHsd (B6, Envigo), Tg(Drd1-cre)EY262Gsat (Drd1-Cre, Gensat), Tg(Adora2a-cre)2MDkde (Adora2A-Cre (Durieux et al., 2009)), Oprm1-2A-Cre (Oprm1-Cre (Märtin et al., 2019)), B6.Cg-Gt(ROSA)26Sor^tm14(CAG-tdTomato)Hze^/J (Ai14, the Jackson Laboratory), B6J.Cg-Gt(ROSA)26Sor^tm95.1(CAG-GCaMP6f)Hze^/MwarJ (Ai95D, the Jackson Laboratory).

### In vitro electrophysiology

#### Slice preparation

Animals were deeply anesthetized using isoflurane before sacrifice by decapitation. Brains were rapidly removed and placed in ice-cold cutting solution. Tilted at 30° horizontal cortico-striatal slices or horizontal slices containing SNpc were prepared on a vibratome (Leica VT 1200S, Leica Biosystems) in ice-cold choline chloride-based cutting solution of the following composition (mM): 117 C_5_H_14_ClNO (choline chloride), 2.5 KCl, 1.25 NaH_2_PO_4_, 7 MgCl_2_, 26 NaHCO_3_, 0.5 CaCl_2_, 15 D-glucose equilibrated with 95% O_2_-5% CO_2_ mixture. Slice thickness varied depending on the age of the animals (300 μm for P7–P21, 220 μm for adults). After preparation, brain slices were allowed to recover for 1 h at 34°C in artificial CSF with Mg^2+^ concentration elevated to 3 mM. Electrophysiological recordings were done at 30° in ACSF (mM: 124 NaCl, 3 KCl, 1.25 NaH_2_PO_4_, 1 MgSO_4_, 26 NaHCO_3_, 2 CaCl_2_, 15 D-glucose) with perfusion speed 1-2ml/min. In cortico-striatal slices, the recording region was dorsal striatum, without distinguishing between dorso-medial and dorso-lateral parts. Cells were visualized with Olympus BX51WI microscope, equipped with LUMPlanFL N 40x/0.8w water immersion objective and Prime BSI Express sCMOS camera (Teledyne Photometrics). CoolLED pE-300 was used as a source of green/blue light. Whole-cell patch-clamp borosilicate-glass pipets (Harvard Apparatus) were obtained with a horizontal two-stage puller (P-1000, Sutter Instruments) and had a resistance typically between 4 and 6 MΩ. Current- and voltage-clamp recordings were obtained with Multiclamp 700A and Digidata 1440A (Molecular Devices) or NI USB-6341 (National Instruments). Online filtering was achieved using the Bessel 10 kHz filtering with a sampling interval of 20 kHz. All cells were first recorded in voltage-clamp mode at holding potential of -70 mV. Series resistance was monitored by injection of 5 mV voltage steps, and if the change in series resistance during the experiment exceeded 30%, the recording was discarded. Either Clampex 10.7 or WinLTP2.32 were used for the acquisition. Data analysis was done with Clampfit10.7, WinLTP2.32(Anderson & Collingridge, 2007) and MiniAnalysis 6.0.3.

#### Intrinsic excitability

RMP and intrinsic excitability of SPNs were recorded in whole-cell configuration and current clamp mode, using filling solution of the following composition (in millimolar): 126 K gluconate, 15 KCl, 5 NaCl, 10 HEPES, 4 MgATP, and 0.5 Na_2_GTP, pH 7.2. RMP was recorded without injecting any background current. AP firing was evoked by current injections, increased in steps of 10 pA, starting from the holding current, needed to keep the cell at -70 mV. Voltage values were not corrected for liquid junction potential, which was 14.4 mV(Marino et al., 2014).

#### Cell-attached recordings

Spontaneous APs were recorded in the cell-attach configuration with glass pipettes filled with ACSF. After obtaining a tight seal, membrane currents were recorded in I=0 mode.

#### mIPSCs

DA neurons in SNpc were selected visually based on the location and large cell bodies. DA identity was then confirmed by the presence of I_h_ and co-staining of biocytin-filled cells with TH in the slices fixed after electrophysiological experiment. Cells were recorded in whole-cell configuration and voltage clamp mode at -60 mV, using filling solution of the following composition (in millimolar): 105 CsCl, 5.3 CaCl_2_, 4 MgCl_2_, 10 HEPES, 10 EGTA, 5 lidocaine *N*-ethyl chloride, 5 Na_2_-phosphocreatine, 4 MgATP, and 0.5 Na_2_GTP, 2% biocytin, pH 7.2. I_h_ was recorded immediately after breaking the seal by applying voltage injections, decreased in steps of 10 mV. mIPSCs were recorded during 10 minutes in the presence of 1 µM voltage-gated sodium channel blocker TTX, 10 µM AMPA receptor blocker NBQX and 1 µM GABA_B_ receptor blocker CGP55845. After recording, brain slices were transferred to 4% PFA-PBS solution and stored overnight at 4°.

### Ca^2+^ imaging

Living cortico-striatal slices were prepared from neonatal D1-Cre/Ai95D and A2a-Cre/Ai95D animals similarly as for electrophysiological recordings. After recovery period, slices were placed to the recording chamber of the electrophysiological setup and perfused with ACSF at a rate of 2 ml/min. Imaging was done with Prime BSI Express camera and open-source imaging application Micro-Manager1.4/ImageJ at 200 ms interval, 2×2 binning and 50 ms exposure time. Baseline, drug application and wash-out were recorded for 1 minute. Imaging analysis was done using ImageJ and custom-made software for MATLAB (Bonifazi et al., 2009). Striosomal region was selected on the maximal projection of the image stack, based on the brighter GCaMP signal. Active cells were selected on the standard deviation projection of the image stack. Frequency of Ca^2+^ events was calculated as the number of events per minute in all active cells of striosomal or matrix region of the slice.

### Viral transfection and CNO injections

Oprm1-Cre animals and wild type littermates received the injection of the viral construct to dorsal striatum at P1, following the method described in (Olivetti et al., 2020) with modifications. Animals were deeply anesthetized by isoflurane and placed into stereotaxic frame with the body temperature controlled. The target coordinates were determined in reference to the vascular lambda (AP 2.4; ML 1.2; DV 2.3 and 2.7). The skin and scull were gently perforated with needle, which was later used to deliver the viral constructs. pAAV-hSyn-DIO-HA-hM4D(Gi)-IRES-mCitrine alone or with pAAV-hSyn-DIO-EGFP were slowly injected into striatum, total injection volume was 150-200 nL per injection point. pAAV-hSyn-DIO-HA-hM4D(Gi)-IRES-mCitrine and pAAV-hSyn-DIO-EGFP were a gift from Bryan Roth (Addgene viral prep # 50455-AAV8, http://n2t.net/addgene:50455, RRID:Addgene_50455; Addgene viral prep # 50457-AAV1, http://n2t.net/addgene:50457, RRID:Addgene_50457). After injection, animals were left to recover on the heating pad and returned to the home cage after that.

Starting from P6 mice were subcutaneously injected, twice a day (9:00 and 18:00) with CNO dissolved in sterile saline solution, 30 mg/kg (clozapine-N-oxide dihydrochloride, cat #HB6149, HelloBio) or sterile saline solution (0.9% NaCl), consecutively for 9 days. At P52-P79 mice were sacrificed for electrophysiology or immunostaining experiments. The administration method was chosen based on the fact that it takes longer for the concentration of small molecules to reach maximal levels in the interstitial brain fluid when they are injected subcutaneously, compared to intraperitoneally injected (Durk et al., 2018; Shoyaib et al., 2019). This approach helps to prolong the effective time of the drug, without sharply reaching the maximal concentration of the drug in the brain. Pharmacokinetics of CNO after subcutaneous injection was estimated in non-human primates only (Raper et al., 2017). Maximal concentration of CNO in cerebrospinal fluid after 10 mg/kg injection reached the values of 120 nM. Based on the facts that EC50 for hM4Di is 8 nM, but the effective concentration of CNO in the slices *in vivo* is 10 µM (Jendryka et al., 2019), we increased the injection dosage three times. CNO treatment of wild-type animals, not expressing hM4Di, did not cause any detectable effects in our dataset (Fig. 6).

### Immunostaining

Biocytin and TH immunostaining was performed on the slices, fixed after patch clamp experiments. MOR, ChAT, mCitrine and gephyrin/TH stainings were done on the free-floating slices collected after transcardial animal perfusion. For that, mice were deeply anesthetized with isoflurane (≥4%), and after confirming the depth of anesthesia by absence of the toe-pinch reflex they were transcardially perfused with 4% paraformaldehyde (PFA) in phosphate buffered saline (PBS) preceded with PBS. Brains were postfixed in 4% PFA overnight at 4 °C, and the next day, coronal sections, 50µm thickness, were cut on vibratome (Leica VT 1200S, Leica Biosystems). Slices were collected in tissue culture plates and washed three times in PBS (10 mins).

#### MOR staining

Slices were incubated in blocking solution (5% normal goat serum, 0.4% Triton-X in PBS) for one hour at room temperature. Slices were then incubated overnight in rabbit-anti-MOR primary antibody (1:1000, ab10275, Abcam) diluted in blocking solution at 4 °C. Next day, slices were washed three times in PBST (PBS, 0.4% Triton X-100) and three times in PBS (10 mins); and then incubated in secondary antibody Alexa Fluor goat anti rabbit 488 (1:500, cat# A-11008, Invitrogen), diluted in blocking solution, for two hours at room temperature, protected from light.

#### ChAT staining

The heat induced antigen retrieval was performed for 15 mins in sodium citrate buffer pH 6.0 in water bath at 80 °C. Next, slices were washed three times in PBS (5mins) and incubated in blocking solution (1% BSA, 10% donkey serum, 0.3% Triton X-100 in PBS) for one hour at room temperature. Goat anti-ChAT primary antibody (1:100, AB144P Abcam) was diluted in blocking solution, and slices were incubated overnight in shaker at 4 °C. Afterwards, slices were washed in PBS three times (5 mins) and incubated in the secondary antibodies Alexa Fluor donkey anti goat 488 (1:500, cat# A-11055, Invitrogen), diluted in blocking solution for two hours at room temperature, protected from light.

#### mCitrine staining

Slices were incubated in blocking solution (5% normal goat serum, 0.4% Triton-X in PBS) for one hour at room temperature. Slices were then incubated overnight in rabbit-anti-GFP primary antibody (1:1000, ab290, Abcam) diluted in blocking solution at 4 °C. Next day, slices were washed three times in PBST (PBS, 0.4% Triton X-100) and three times in PBS (10 mins); and then incubated in secondary antibody Alexa Fluor goat anti rabbit 568 (1:1000, cat# A-11011, Invitrogen), diluted in blocking solution, for two hours at room temperature, protected from light.

#### Biocytin and TH staining

Patched slices were washed and permeabilized three times (10 mins) in PBST (PBS, 0.4% Triton X-100). Next, slices were incubated in blocking solution (0,4% Triton X-100, 5% normal goat serum in PBS) for one hour at room temperature. Slices were then incubated overnight in primary antibody Anti-Tyrosine Hydroxylase Antibody from rabbit (1:500, cat# AB152, Sigma Aldrich) diluted in blocking solution at 4 °C. Next day, slices were washed three times in PBS (10 mins) and then incubated in secondary antibodies Alexa Fluor goat anti rabbit 568 (1:1000, cat# A-11011, Invitrogen) and streptavidin Alexa fluor 488 (1:500 Streptavidin, cat# S32354, Invitrogen) diluted in blocking solution, for two and a half hours at room temperature, protected from light.

#### Gephyrin and TH staining

The heat induced antigen retrieval was performed for 20 mins in sodium citrate buffer pH 6.0 in water bath at 80 °C. Next, slices were washed three times in PBS (5mins) and incubated in blocking solution (10% normal goat serum, 0.2% Triton X-100 in PBS) for one hour at room temperature. Primary antibodies Anti-Gephyrin Monoclonal Mouse Antibody (1:250, cat# 147 021, Synaptic Systems) and Anti-Tyrosine Hydroxylase Antibody from rabbit (1:500, cat# AB152, Sigma Aldrich) were diluted in blocking solution, and slices were incubated over night at 4 °C. Afterwards, slices were washed in PBS three times (10 mins) and incubated in the secondary antibodies Alexa Fluor goat anti rabbit 568 (1:1000, cat# A-11011, Invitrogen) and Alexa Fluor goat anti mouse 647 (1:1000, cat# A-21236, Invitrogen) diluted in blocking solution for two hours at room temperature, protected from light.

### Imaging

After immunostaining, slices were washed three times in PBS (10 mins; for MOR staining six times by 10 min) and mounted on super frost plus adhesion microscope slides with Fluoromount-G mounting medium (cat # 00-4958-02, Invitrogen). Before imaging, slides were left to dry overnight and afterwards were stored at 4 °C, protected from the light.

For analysis of the **dendritic tree** of patched dopaminergic neurons in substantia nigra pars compacta imaging was done with Leica Upright SP8 confocal microscope with Blue laser (488nm) and HC PL APO 20x/0.75 IMM (immersion) objective. To ensure that the entire dendritic tree was imaged the tile, z-stack imaging of the patched THC positive (dopaminergic) neuron was done, with z=2 µm and resolution x, y=0.56 µm (1024×1024 pixels). Line average was set to three. Sholl analysis of dendritic arborization for four different experimental groups was done in Image J, with radius set to 10 µm.

For MOR, Chat, and mCitrine **immunostainings**, images were obtained with ApoTome.2, Zeiss microscope. For MOR and Chat immunostainings Plan-APOCHROMAT 10x/045 air objective was used, and red (emission 581 nm) and green (emission 517 nm) filters were used for tdTomato and µ-opioid receptor/Chat stainings, respectively. mCitrine immunostaining was imaged with EC PLAN-NEOFLUAR 5x/016 air objective, with filters red (emission 581 nm) for mCitrine and blue (emission 465 nm) for DAPI signal. Resolution for all images was 2048×2048 pixels.

For **tracing experiment** images were acquired with Leica Upright SP8 confocal microscope, with HC PL APO 63x/1.30 GLYC (glycerol) objective. Laser used were Blue (488 nm) for GFP signal, Red (638 nm) for THC signal and Lime (552 nm) for gephyrin. Representative striosome dendron bouquets images were taken with z stack z= 2 µm and resolution x, y= 0.24 µm (1024×1024 pixels), zoom=0.75. For analysis of synaptic density and relative GFP volume, pinhole size and laser power were set and were same across the samples, while photomultiplier gain was adjusted individually for every sample imaged, ensuring the signal was not over exposed. Step size and pixel size were automatically optimized to 0.33µm (z-stack) and x, y=0.076 µm (608 × 608 pixels) respectively. TH positive (dopaminergic) neuron was randomly selected in substantia nigra pars compacta region and zoomed on (zoom=4), afterwards the z stack was taken. Line average was set to three in all imaging.

Analysis of the density of striosomal projections in SNpc was done using Imaris 9.9.1. The GFP relative volume was calculated as the total green signal volume in voxels per total voxel volume of the image stack, while the relative volume of gephyrin was calculated as total number of dots per total voxel volume of the image stack. Diameter and quality filter of the dots were optimized and set to constant value across the images, while absolutely intensity threshold of green signal was adjusted for each image individually.

## Figures

**Figure S1-related to Fig. 2.**
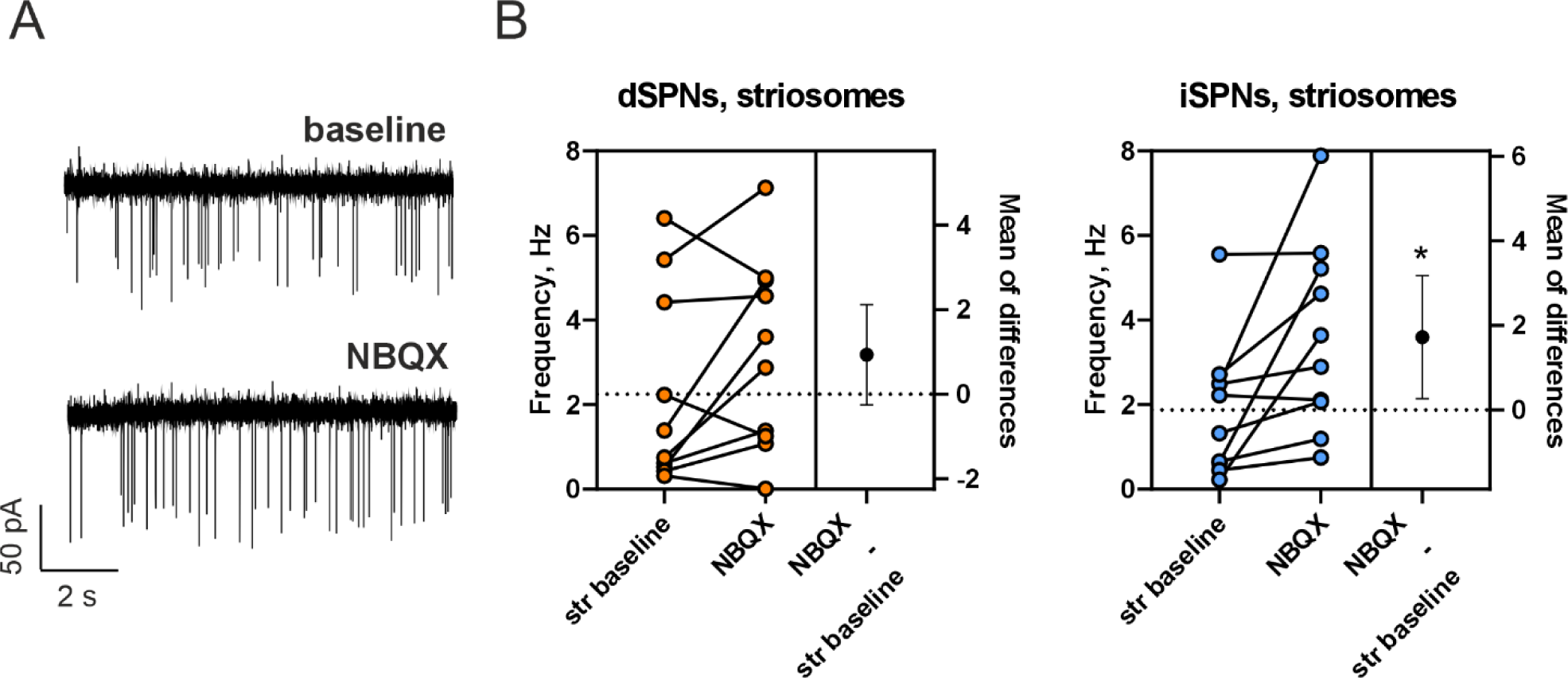
Blockade of AMPA receptors does not suppress spontaneous neonatal activity. A. Example traces of the AP firing, recorded in the cell-attached mode from the striosomal iSPN of the P7 A_2A_-Cre/Ai14 animal, during baseline and NBQX (20 µM) application. B. Quantification of the AP frequency before and during application of NBQX in neonatal (P4-7) dSPNs and iSPNs. Data are presented as individual values and mean of the difference with 95% CI. dSPNs: 10 cells from 4 animals; iSPNSs: 10 cells from 5 animals. *p<0.05, paired t-test.

**Figure S2 – related to Figure 6.**
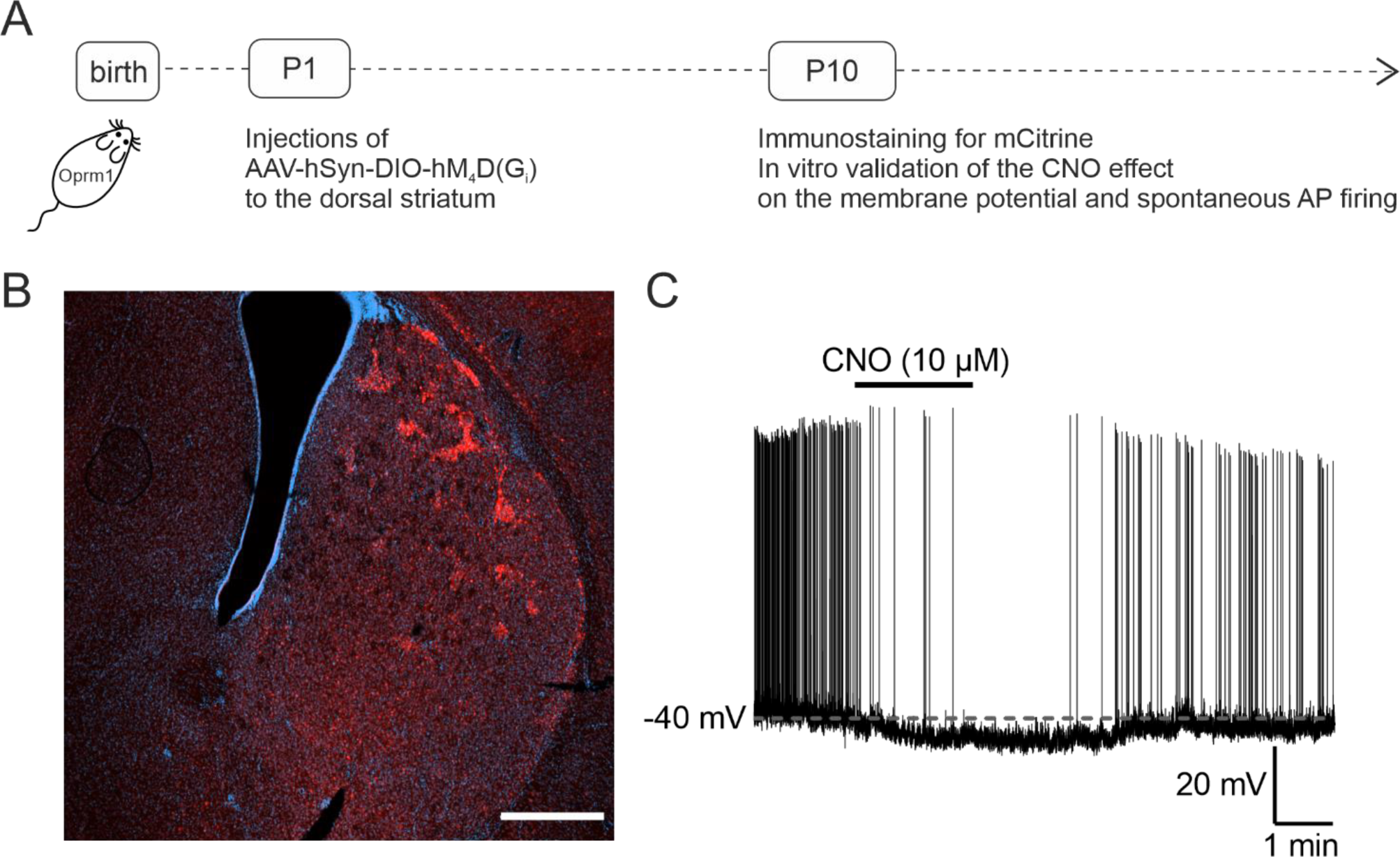
Characterization of the hM_4_D(G_i_) expression at P10. A. Schematic illustration of the experimental timeline. Oprm1-Cre animals received microinjection of AAV-hSyn-DIO-hM_4_D(G_i_) to the dorsal striatum at P1. At P10 animals were sacrificed and used for the immunochemistry or patch-clamp. B. Coronal slice of dorsal striatum with the expression of mCitrine in striosomes. Red: mCitrine staining; blue: DAPI. Scale bar 500 µm. C. Example trace of membrane potential recording of the striosomal SPN during application of 10 µM CNO.

**Figure S3 – related to Figure 6.**
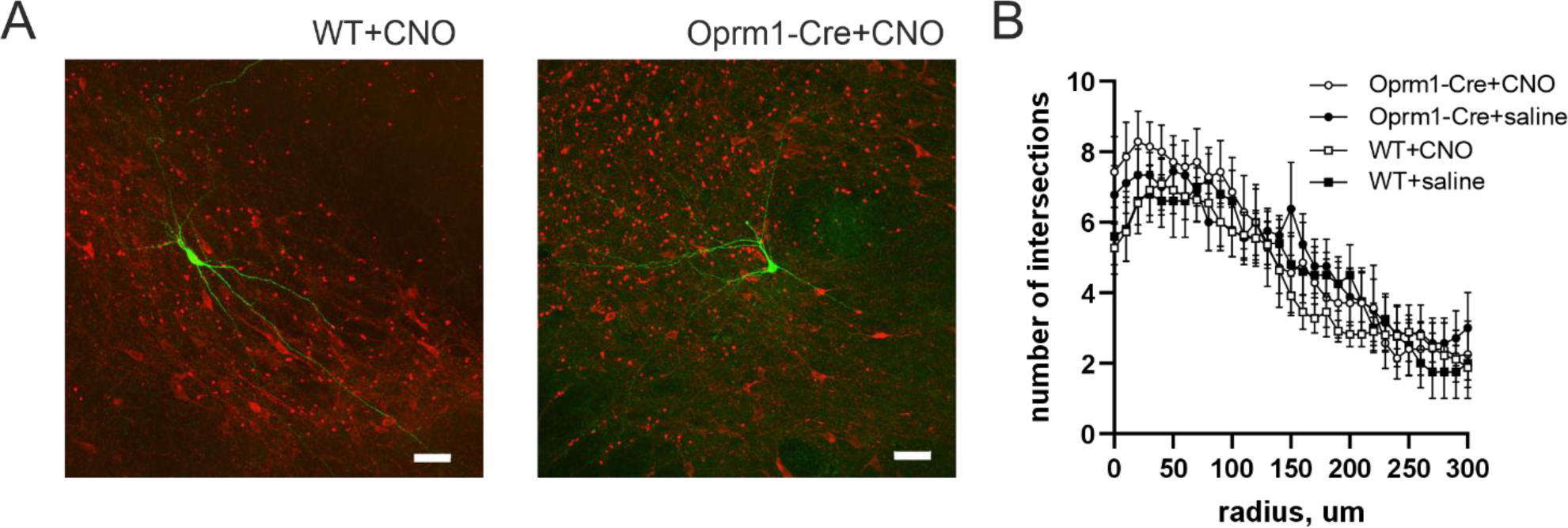
The dendritic tree of DA neurons was not affected by chemogenetic suppression of the neonatal striosomal activity. Oprm1-Cre animals and wild-type littermates received cranial injection of AAV-hSyn-DIO-hM_4_D(G_i_) into dorsal striatum at P1. At P6-P14 animals received s.c. injections of CNO twice a day. Adult animals were used for whole-cell patch clamp experiments, where DA neurons in SNpc were filled by biocytin for the analysis of the dendritic tree. A. Example images of DA cells filled by biocytin from SNpc of adult wild-type and Oprm1-Cre animals after viral transfection and neonatal injections of CNO. B. Sholl analysis graph showing the number of intersections of the dendrites with different radiuses, starting from the soma. Sample size, cells/animals: WT CNO 11/8, Oprm1-Cre CNO 7/5, WT saline 5/3, Oprm1-Cre saline 9/6. Data are presented as mean ± SEM.

## Key resources table

| Reagent or Resource | Source | Identifier |
| --- | --- | --- |
| **Experimental Models: Organisms/Strains** |  |  |
| C57BL/6JRccHsd | Envigo | MGI:6156915 |
| Tg(Drd1-cre)EY262Gsat (D1-Cre) | GENSAT project | MGI:3836631 |
| Tg(Adora2a-cre)2MDkde (A2A-Cre) | Alban de Kerhove d'Exaerde | MGI:3852493 |
| Oprm1-2A-Cre (Oprm1-Cre) | Konstantinos Meletis | own breeding |
| B6.Cg-Gt(ROSA)26Sor<sup>tm14</sup>(CAG-tdTomato)<sup>Hze</sup>/J (Ai14) | the Jackson Laboratory | JAX 007914 |
| B6J.Cg-Gt(ROSA)26Sor<sup>tm95.1</sup>(CAG-GCaMP6f)<sup>Hze</sup>/MwarJ (Ai95D) | the Jackson Laboratory | JAX 028865 |
| **Viral constructs** |  |  |
| pAAV-hSyn-DIO-EGFP | Bryan Roth, Addgene | 50457-AAV1 |
| pAAV-hSyn-DIO-HA-hM4D(Gi)-IRES-mCitrine | Bryan Roth, Addgene | 50455-AAV8 |
| **Antibodies** |  |  |
| rabbit-anti-MOR | Abcam | ab10275 |
| Goat anti-ChAT | Abcam | AB144P |
| rabbit-anti-GFP | Abcam | ab290 |
| rabbit-anti-TH | Merck Sigma Aldrich | AB152 |
| mouse-anti-gephyrin | Synaptic Systems | 147 021 |
| goat anti rabbit 488 | Invitrogen | A-11008 |
| donkey anti goat 488 | Invitrogen | A-11055 |
| goat anti rabbit 568 | Invitrogen | A-11011 |
| streptavidin Alexa fluor 488 | Invitrogen | S32354 |
| goat anti mouse 647 | Invitrogen | A-21236 |
| **Chemicals** |  |  |
| Tetrodotoxin citrate (TTX) | HelloBio | HB1035 |
| NBQX disodium salt | HelloBio | HB0443 |
| CGP55845 hydrochloride | HelloBio | HB0960 |
| biocytin | Merck Sigma Aldrich | B4261 |
| VU0255035 | Tocris/Bio-Techne | 3727/5 |
| clozapine-N-oxide dihydrochloride (CNO) | HelloBio | HB6149 |

